# Cerebrospinal fluid formation is controlled by membrane transporters to modulate intracranial pressure

**DOI:** 10.1101/2021.12.10.472067

**Authors:** Eva K. Oernbo, Annette B. Steffensen, Pooya Razzaghi Khamesi, Trine L. Toft-Bertelsen, Dagne Barbuskaite, Frederik Vilhardt, Niklas J. Gerkau, Katerina Tritsaris, Anja H. Simonsen, Sara D. Lolansen, Søren N. Andreassen, Steen G. Hasselbalch, Thomas Zeuthen, Christine R. Rose, Vartan Kurtcuoglu, Nanna MacAulay

**Author notes:** These authors contributed equally to this work (Eva K. Oernbo, Annette B. Steffensen). Correspondence: Nanna MacAulay, University of Copenhagen, Faculty of Health and Medical Sciences, Department of Neuroscience, Blegdamsvej 3, DK-2200 Copenhagen, Denmark.

## Abstract

Disturbances in the brain fluid balance can lead to life-threatening elevation in the intracranial pressure (ICP), which represents a vast clinical challenge. Nevertheless, the molecular mechanisms governing cerebrospinal fluid (CSF) secretion are largely unresolved, thus preventing targeted and efficient pharmaceutical therapy of cerebral pathologies involving elevated ICP. Here, we employed experimental rats to demonstrate low osmotic water permeability of the choroid plexus, lack of an osmotic gradient across this tissue, and robust CSF secretion against osmotic gradients. Together, these results illustrate that CSF secretion occurs independently of conventional osmosis, which challenges the existing assumption that CSF production is driven entirely by bulk osmotic forces across the CSF-secreting choroid plexus. Instead, we reveal that the choroidal Na^+^/K^+^/Cl^−^ cotransporter NKCC1, Na^+^/HCO_3_^−^ cotransporter NBCe2, and Na^+^/K^+^-ATPase are actively involved in CSF production and propose a molecular mode of water transport supporting CSF production in this secretory tissue. Further, we demonstrate that inhibition of NKCC1 directly reduces the ICP, illustrating that altered CSF secretion may be employed as a strategy to modulate ICP. These insights identify new promising therapeutic targets against brain pathologies associated with elevated ICP.

## Introduction

Our brain contains 80% water in the form of cerebrospinal fluid (CSF) that surrounds the brain tissue and fills ventricles and interstitial spaces. The CSF protects the brain from mechanical insults and serves as a route for dispersion of hormones, nutrients, and metabolites between brain structures and neighboring cells. The CSF is replenished at a rate of approximately 500 ml/day in the adult human ^1^, the majority of which is secreted across the choroid plexus epithelium residing in the ventricles ^2,3^. The CSF disperses in the brain prior to entering the proposed exit routes along the lymphatic and/or perineural pathways ^4–7^. CSF dynamics are finely tuned to ensure stable brain water content and an intracranial pressure (ICP) within the physiological range ^8^. However, many brain pathologies are associated with CSF accumulation, e.g., in the form of hydrocephalus or brain edema. An elevation in ICP may be debilitating or even fatal to the patient if left untreated. The prevalent mode of ICP management under these conditions relies on surgical intervention such as ventricular shunt implantation or craniectomy, which are highly invasive and associated with severe side effects and/or need for recurring revisions ^9,10^. Pharmacological approaches towards ICP reduction generally fail due to questionable efficiency and/or intolerable side effects ^11–13^. This failure relates to our incomplete understanding of the molecular mechanisms underlying the CSF secretion and therefore the lack of rational targets for directed and specific therapy.

It is generally assumed that CSF secretion takes place by conventional osmotic water transport, where water follows the osmotic gradient that results from electrolyte transport across the choroid plexus epithelium ^14^. For such manner of osmotic water flux to sustain the entire CSF secretion, however, a sizeable osmotic gradient and sufficient water permeability must exist across the choroidal epithelium.

Alternatively, local osmotic gradients could potentially generate favorable conditions for fluid flow across the choroid plexus epithelium. Neither of these have been demonstrated experimentally, rendering the molecular mechanism of CSF secretion unresolved ^3^. The Na^+^/K^+^/2Cl^−^ cotransporter 1 (NKCC1) was recently proposed as a key contributor to CSF secretion in mice ^15^ and rats ^16^, and may do so via its ability to promote fluid transport ^17,18^. This manner of water transport appears to occur by a mechanism within the protein itself ^19,20^ and therefore does not require a bulk trans-epithelial osmotic gradient ^21^. Other transporters expressed in the choroid plexus epithelial cells may share this feature.

In this study, we aimed to resolve the pivotal question of how CSF is secreted across the choroid plexus, and demonstrate that CSF secretion can occur independently of a bulk trans-epithelial osmotic gradient. The CSF secretion across the luminal membrane does not depend on local osmotic gradients, but is supported by a concerted effort of cotransporters and pumps present in the choroidal epithelium, possibly via transport of water by a mechanism inherent in the transport proteins. Such mode of fluid transport eliminates the necessity for large osmotic gradients between the vasculature and the brain. Importantly, the definition of such specific molecular targets will allow pharmacological interference with the CSF secretion to reduce ICP in disease.

## Methods

### Patient samples

Fourteen elderly and evenly sex-distributed individuals (mean age: 67 y, range: 58 - 81 y, 9 F / 11 M) were included in the study. Lumbar CSF samples and corresponding serum samples were obtained from the Danish Dementia Biobank at the Danish Dementia Research Centre (Copenhagen University Hospital, Rigshospitalet, DK) together with relevant clinical information. Patients gave consent for using samples for research purposes according to the guidelines given by the Ethics committee of the Capital Region of Denmark. All elderly control individuals were screened, but cleared, for dementia, mild cognitive impairment, and normal pressure hydrocephalus ^22–24^. To evaluate the impact of different blood sampling methods on the osmolality (in mosmol per kg, termed mOsm henceforward), blood samples from seven healthy volunteers were included (mean age: 32 y, range: 23-39 y, 5 F / 2 M).

### Animals

All animal experiments conformed to the legislations for animal protection and care in the European Community Council Directive (2010/63/EU) and followed all ethical regulations under the animal permission no. 2016-15-0201-00944 and 2018-15-0201-01595 authorized by the Danish Animal Inspectorate or the institutional act no. O50/05 approved by the Animal Welfare Office at the Animal Care and Use Facility of the Heinrich Heine University Düsseldorf. Adult male Sprague Dawley rats (Janvier Labs) at 9 ± 1 weeks of age were used for physiological experimentation. The animals were housed with a 12:12 light cycle and had free access to water and food. Animals were randomly allocated to each treatment group, and all experimental work was performed and reported in compliance with the ARRIVE guidelines ^25^. The sample size for all animal experiments was determined with power calculations ^15^. Additional biosamples (choroid plexus, CSF, blood) were obtained from adult female Danish landrace pigs (weight ∼ kg, conventional producer with blue SPF status) under the animal permission 2016-15-0201-00845.

### Anesthesia

Rats were anesthetized with an intraperitoneal (i.p.) injection of xylazine and ketamine (6 mg/ml and 60 mg/ml in sterile water, 0.17 ml per 100 g body weight, ScanVet). To sustain anesthesia, rats were re-dosed with a half ketamine dose upon detection of foot reflexes after pinching (approximately every 20-40 min). When indicated, isoflurane (Attane vet, 1000 mg/g isoflurane, ScanVet) was employed using 4 - 5% (mixed with 0.9 l min^−1^ air / 0.1 l min^−1^ O_2_ for terminal procedures or 1.8 l min^−1^ air / 0.1 l min^−1^ O_2_ for survival procedures) to induce anesthesia in an induction chamber and 2 - 2.5% to maintain anesthesia through a face mask, which was gradually decreased to 1% during the surgery.

### Solutions and chemicals

The majority of the experiments were conducted in CO_2_/HCO_3_-buffered artificial cerebrospinal fluid (HCO_3_-aCSF; (in mM) 120 NaCl, 2.5 KCl, 2.5 CaCl_2_, 1.3 MgSO_4_, 1 NaH_2_PO_4_, 10 glucose, 25 NaHCO_3_, bubbled with 95% O_2_/5% CO_2_ to obtain a pH of 7.4). In experiments where the solution could not be equilibrated with 95% O_2_/5% CO_2,_ it was instead buffered by HEPES (HEPES-aCSF; (in mM) 120 NaCl, 2.5 KCl, 2.5 CaCl_2_, 1.3 MgSO_4_, 1 NaH_2_PO_4_, 10 glucose, 17 Na-HEPES, adjusted to pH 7.4 with NaOH). All pharmacological inhibitors were purchased from Sigma. Pharmacological inhibitors were dissolved in DMSO and kept as stock solutions at −20°C (bumetanide: B3023, cariporide: SML1360, S0859: SML0638) or dissolved directly into the solution (furosemide: F4381, ouabain: O3125, DIDS: D3514) on the day of the experiment. All solutions included the appropriate vehicle; mostly addition of 0.05% DMSO (D8418, Sigma). In a subset of experiments containing higher DMSO concentration, the osmolality of the solution was maintained at 302 ± 1 mOsm by reducing the [NaCl] (e.g. to 108 mM with 0.25% DMSO). The fluorescent dyes calcein-AM (C1359, Sigma or C1430, ThermoFisherScientific), BCECF-AM (2’,7’-bis-(2-carboxyethyl)-5-(and-6)-carboxyfluorescein, acetoxymethyl ester, 4011E, ION Biosciences, dissolved in 20 % Fluronic, F127), and SBFI-AM (sodium-binding benzofuran isophthalate acetoxymethyol ester, 2021E, ION Biosciences, dissolved in 20 % Pluronic, F127) were stored in DMSO stock at −20°C, while TRITC-dextran (tetramethylrhodamine isothiocyanate-dextran, MW = 150,000; T1287, Sigma), and carboxylate (MW=1,091, IRDye 800 CW, P/N 929-08972, LI-COR Biosciences) were dissolved directly in aCSF.

### Physiological parameters of anesthetized rats

Rats were fully anesthetized during all *in vivo* experiments and their body temperature was maintained at 37 °C by a homeothermic monitoring system (Harvard Apparatus). For anesthetic protocols lasting more than 30 min, mechanical ventilation was included to ensure stable respiratory partial pressure of carbon dioxide and arterial oxygen saturation and thus stable plasma pH and electrolyte content. A surgical tracheotomy was performed and the ventilation controlled by the VentElite system (Harvard Apparatus) by 0.9 l min^−1^ humidified air mixed with 0.1 l min^−1^ O_2_ adjusted with approximately 3 ml per breath, 80 breath min^−1^, a Positive End-Expiratory Pressure (PEEP) at 2 cm, and 10 % sight for a ∼400 g rat. The ventilation settings were optimized for each animal using a capnograph (Type 340, Harvard Apparatus) and a pulse oximeter (MouseOx® Plus, Starr Life Sciences) after system calibration with respiratory pCO_2_ (4.5 - 5 kPa) and pO_2_ (13.3 - 17.3 kPa) and arterial oxygen saturation (98.8 - 99.4 %) (ABL90, Radiometer). Blood pressure measurements were obtained by introducing a heparinized saline-filled catheter (15 U heparin / ml in 0.9% NaCl) into the femoral artery connected to a transducer (APT300, Hugo Sachs Elektronik). The mean arterial blood pressure and heart rate were collected by the HSE Pressure Measurement System and BDAS v2.0 software (Huge Sachs Elektronik) (See Supplementary Fig. 1a-b). Blood electrolytes (Na^+^, K^+^, Cl^−^) and the hemoglobin (Hb) content were determined using an ABL80 (Radiometer, Supplementary Fig. 1f-i).

### Ventriculo-cisternal perfusion

Anaesthetized and ventilated rats were placed in a stereotaxic frame (Harvard Apparatus) in a prone position. A dorsal midline incision was made over the skull and upper cervical spine to expose the cranium, after which a burr hole was drilled (0.5 mm drill bit) using the coordinates: 1.3 mm posterior to bregma, 1.8 mm lateral to the midline, and ∼0.6 mm ventral through the skull. A brain infusion cannula (Brain infusion kit2 with 1 mm adjustment spacers, Alzet) was placed into the lateral ventricle through the burr hole and glued to the skull (Superglue, Pelikan). The perfusion solution (HCO_3_^−^-aCSF containing 1 mg/ml TRITC-dextran and 0.05 - 0.25% DMSO vehicle) was pH calibrated and heated to 37 °C by an inline heater (SF-28, Warner Instruments) prior to entering the infusion cannula. After securing access through the neck muscle layer, a glass capillary (30-0067, Harvard Apparatus pulled by a Brown Micropipette puller, Model P-97, Sutter Instruments) was introduced at a 5° angle into cisterna magna (7.5 mm distal to the occipital bone and 1.5 mm lateral to the muscle-midline). The HCO_3_^−^-aCSF was continuously infused into the lateral ventricle at a rate of 9 µl min^−1^ using a peristaltic pump. CSF was collected from cisterna magna in 5 min intervals from a second glass capillary (30-0065, Havard Apparatus) placed into the first capillary. For experiments including pharmacological inhibitors, the solution was changed after 1 h infusion to a HCO_3_^−^-aCSF containing the transport inhibitor (100 µM bumetanide, 5 mM ouabain, 1.5 mM DIDS, or 50 µM cariporide, all expected to be diluted at least ∼1:1 upon ventricular entrance, due to the comparable rates of aCSF delivery and CSF secretion). The fluorescence in the collected sample fluids was measured using the Synery^TM^ Neo2 Multi-mode Microplate Reader (BioTek Instruments). The CSF production rate was calculated from the equation:

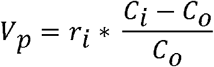

where *V_p_* = CSF production rate (µl min^−1^), *r_i_* = infusion rate (µl min^−1^), *C_i_* = fluorescence of inflow solution, *C_o_* = fluorescence of outflow solution, calculated 50-65 min after infusion initiation (baseline) and 100-120 min after infusion initiation (inhibitor effect). Different ventricular osmolalities were obtained by unilateral infusion of a hyper- or hyposmolar HCO_3_^−^-aCSF. Here, the control aCSF solution (307 mOsm) contained 100 mOsm less NaCl, which was replaced with equiosmolar mannitol (105 mM). In this manner, a hyposmolar solution could be obtained by removal of mannitol (and the hyperosmolar solution with addition of mannitol). Thereby, only the osmolality, and not the electrolyte content, was altered during the experimental procedure. In one experimental series, the aCSF osmolality was adjusted with NaCl (55 mM) instead of mannitol to mimic a physiological solution. With the continued CSF production and the pump infusion rate, we estimate a 1:1 dilution of the infused aCSF with that of the existing CSF. The unilateral infusion of aCSF of a different osmolality was initiated after cisterna magna puncturing and continued for 75 min before termination. The CSF production rate was calculated 50-75 min after infusion initiation. In a subset of experiments, blood pressure measurements from the femoral artery were obtained. All experiments were randomized, and data analyses performed under blinded conditions.

### Live imaging of CSF tracer

Anesthetized rats were placed in a stereotaxic frame and a burr hole drilled at coordinates 1.3 mm posterior to bregma and 1.8 mm lateral to the midline. Ventricular injections were performed with a hamilton syringe (RN 0.40, G27, a20, Agntho’s) inserted 4 mm into the right ventricle. 15 μl HCO_3_^−^-aCSF containing vehicle (0.1% DMSO) or each transport inhibitor (200 μM bumetanide, 2 mM ouabain, 100 μM cariporide or 300 μM S0859) was injected over 10 s. These inhibitors are expected to be swiftly diluted (approximately 10-fold) in the native ∼ 150 μl CSF in both the ipsi- and contralateral ventricle, which leads to estimated functional ventricular concentrations of 20 μM bumetanide, 200 μM ouabain, 30 μM S0859 or 10 μM cariporide. HCO_3_^−^- aCSF containing carboxylate dye (10 μM) including the tested inhibitor was injected 5 min later. The rat was swiftly transferred to the Pearl Trilogy Small Animal Imaging System (LI-COR) and the head fastened in a custom-made tooth-holder that enabled stable head position during imaging. Exactly 1 min after the second injection, imaging was initiated with 30 s intervals for 5 min (800 nm channel, 85 μm resolution). A white image was obtained at the termination of each experiment. To verify bilateral reach of inhibitor and carboxylate, the ventricles of separated hemispheres from the isolated rat brain were imaged subsequently. The fluorescence signals were analyzed in Image Studio 5.2 with the ROIs defined from the white image as a square starting at lambda. Quantification of the relative 2D movement of dye over time is shown as fluorescent signals at given time points normalized to the first image fluorescent signal. Quantifications were performed in a blinded manner.

### ICP monitoring

Anesthetized and ventilated rats, kept at 37 °C body temperature, were placed into a stereotactic frame. The skull was exposed through a dorsal midline incision, a burr hole drilled, and a 4 mm brain infusion cannula was placed into the lateral ventricle (coordinates of 1.3 mm posterior to bregma, 1.8 mm lateral to the midline, and ∼0.6 mm ventral through the skull). A cranial window (∼ 4 mm diameter) was drilled on the contralateral side of the skull with care taken not to damage the dura. An epidural probe (PlasticsOne, C313G, with the ICP cannula cut to be level with the base of the pedestal) was lowered into the hole, and secured with dental resin cement (Panavia SA Cement, Kuraray Noritake Dental Inc.). The ICP cannula was pre-filled with HEPES-aCSF and connected to a pressure transducer (APT300) and transducer amplifier module TAM-A (both from Hugo Sachs Elektronik). The pressure signal was visualized and recorded with a 1 kHz sampling rate using BDAS Basic Data Acquisition Software (Harvard Apparatus, Hugo Sachs Elektronik). To assure a continuous fluid column between the dura and the epidural probe, approximately 5 µl of HEPES-aCSF was injected through the epidural probe, after which the ICP signal stabilized within 15 minutes. Ventricular infusion of heated and pH-equilibrated HCO_3_^−^-aCSF with vehicle (0.9 % DMSO) was initiated with 0.5 µl/min by a peristaltic pump (or 1.5 µl/min for a selected set of experiments) for 45 min with subsequent solution switch to an HCO_3_^−^-aCSF containing 1.8 mM bumetanide (expected dilution to an approximate ventricular concentration of 50 µM, based on the ventricular CSF volume and the slow infusion rate of the inhibitor) and continued for 100 minutes. Jugular compression was applied at the initiation and termination of the experiment to confirm proper ICP recording.

### Telemetric pressure monitoring

Telemetric ICP monitoring was performed using KAHA Sciences rat dual pressure telemetric system. Animals were anaesthetized by isoflurane inhalation, the body temperature was maintained at 37 °C, and the surgical areas sterilized using 0.5 % chlorhexidine (Medic). Analgesia was injected s.c. prior and post-surgery (48 h) and contained 5 mg/kg Caprofen (Norodyl Vet, Norbrook), 0.05 mg/kg buprenorphine (Tamgesic, Indivior), and 200 + 40 mg/kg sulfadiazin and trimethoprim (Borgal Vet). The implantation procedure was performed under aseptic conditions as described in ^26^. Briefly, the blood flow of the abdominal aorta was occluded (4-0 silk suture threads, Vömel) and the tip of the pressure probe inserted into the aorta using a bent 23G needle. The hole was sealed with tissue adhesive (Histoacryl, Enbucrilate; B. Braun) and gluing of a square (5 × 5 mm) of surgical mesh on to the area (PETKM2002, SurgicalMesh; Textile Development Associates). The telemetric device was secured to the abdominal wall and the abdominal muscles sutured (with 4-0 absorbable Vicryl suture, Ethicon), leaving the ICP second pressure probe protruding. The pressure probe was tunneled towards the base of the skull with a 6 mm diameter stainless steel straw (Ecostrawzand) and the abdominal incision closed with wound clips (11 × 2 mm Michel clips, B. Braun). The animal was placed into the stereotactic frame (Harvard apparatus), its skull exposed prior to drilling two holes on the ipsilateral side of the skull (one with a 1.2 mm burr bit and one further anterior with a 1.4 mm burr bit). A stainless-steel screw (00-96×3/32, Bilaney Consultants GmbH) was inserted in the former hole to anchor the pressure probe, which is placed in the latter hole. The pressure probe was sealed with spongostan (Ethicon) and secured with surgical mesh and tissue adhesive. Dental impression material (Take 1 Advanced) was applied over the catheter and the screw and the skin incision was closed with non-absorbable 4-0 EthilonII suture (Ethicon). Telemetric recordings were initiated 9 days after the implantation by placing the animal cages on the TR181 Smart Pads (Kaha Sciences). The telemetric signals were sampled at 1 kHz with PowerLab and LabChart software (v8.0, ADInstruments) and continued for 3-4 days to obtain the baseline ICP, mean arterial pressure, and heart rate.

### CSF and blood osmolality

The osmolality of CSF and blood was determined in rats, pigs, and human samples. In animals, the CSF was extracted by cisterna magna puncture at the atlanto-occipital membrane immediately after anesthesia induction and stored in 2 ml polypropylene tubes (62.610.018, Hounisen) ^27^. A mix of arterial and venous blood was extracted from the neck region of rats after decapitation, while venous blood was collected directly from the heart or ear of the pigs. In humans, CSF samples were obtained by lumbar puncture and collected in polypropylene tubes (62.610.018, Hounisen) and blood samples obtained by venipuncture. Several blood collection tubes were used for blood analysis; heparin coated tubes (290353 Kruuse or 454285, Greiner-Bio One), EDTA coated tubes (41.1395.105, Hounisen or 456043, Greiner Bio-One) and serum tubes (45 4067, Geiner-Bio One). Immediately after collection, blood and CSF samples were centrifuged at 2000 g for 10 min at 4 °C (blood collected in serum tubes was allowed to clot for 30 min at room temperature before centrifugation). Pellets were discarded and supernatant of human samples stored in −80 °C pending analysis, while the animal samples were analyzed freshly. Prior to osmolality measurements, animal C (up to max 1 h), while frozen human samples were thawed and kept on ice (max 4 h). 100 μl supernatant was transferred to osmometer tubes (72.690.001, VWR) and the osmolality was determined on duplicate samples using a freezing point depression osmometer (Löser, Type 15) with a ± 1 mOsm accuracy.

### Electron microscopy

Acutely isolated choroid plexus was fixed immediately in 2% glutaraldehyde in phosphate buffer, pH 7.2, before standard processing for either conventional transmission electron microscopy (TEM) or for scanning electron microscopy (SEM). 50 nm TEM sections were examined in a CM 100 microscope and images acquired with an Olympus Veleta camera at a resolution of 2048 × 2048. Scanning EM samples were observed in a FEI Quanta 3D FEG dual beam scanning microscope and images acquired at a resolution of 2048 × 2048. Microvillus length was estimated by the track length function of ImageJ using scanning EM images (to unequivocally follow the same microvillus; n=10 cells, 45 microvilli). The microvilli diameter was determined in TEM cross sections, and the linear density of microvilli was estimated by counting microvilli continuous with the cell surface per μm of bulk membrane (ImageJ); to obtain microvillar density, the result was integrated over 1 μm^2^.

### Mathematical modelling of CSF secretion

A mathematical model of osmotic water transfer into the inter-microvillar space of the choroid plexus luminal membrane was developed based on data from above-described physiological experiments and transmission electron microscopy (see Supplementary Data). The functional unit of the model is the volume between four microvilli, simplified as a hydraulically equivalent circular cylindrical space. Solutes are continuously supplied into this space through the adjacent membrane segments, producing an osmotic gradient that draws intracellular water. Assuming steady state conditions, the local solute flux corresponds to the rate of solute removal by bulk CSF drainage divided by the total lateral and base surface areas of all functional units. Fluid velocity and solute concentration distribution along the longitudinal axis of a functional unit were determined through a set of coupled steady-state differential equations originally proposed by Diamond & Bossert ^28^, see Supplementary Data. The model equations were solved in MATLAB Release 2020b (MathWorks, Natick, MA, USA) using the boundary value problem solver *bvp4c*. The implementation was validated against published results ^28^. Computational mesh independence was tested for and confirmed.

### Volume recordings

Acutely isolated rat choroid plexus was mounted on coverslips coated with Cell-Tak® (Corning) and placed in a POCmini2 open perfusion chamber (PeCon) mounted in a 37 °C heating chamber of an inverted confocal microscope (Carl Zeiss, Cell-Observer with a Yokogawa X1 spinning disk, 10x 0.3 NA EC Plan Neofluar objective). The choroid plexus was loaded with calcein-AM (17 µM, 10 min) prior to washout of excess fluorophore with ∼1 ml/min continuous HEPES-aCSF flow induced by gravity. Imaging was initiated (488 nm, 50 ms exposure) and the emitted fluorescence recorded at an acquisition rate of 1 Hz (Hamamatsu Orca Fusion) with the ZEN software. An additional 100 ± 1 mOsm was added to the HEPES-aCSF by inclusion of either 100 mM mannitol, 55 mM NaCl or 55 mM KCl. All experiments were randomized prior to execution and the subsequent analysis carried out in an experimenter-blinded fashion. Images were converted to black/white (background/choroid plexus) using Gaussian blur threshold in ImageJ (NIH) and the response recorded as the average of the five regions of interest placed with 50 % choroid plexus and 50 % background. The two-dimensional distribution change was employed as a proxy for choroid plexus volume changes and was determined as a function of time.

### Imaging of Na^+^ and pH

Acutely isolated lateral choroid plexus kept in HCO_3_^−^-aCSF was loaded with the Na^+^-sensitive fluorescent probe SBFI-AM (150 µM) or with the pH-sensitive fluorescent probe BCECF-AM (125 µM) by pressure-application onto several regions of the isolated tissue. After dye loading, the tissue was washed by HCO_3_^−^-aCSF perfusion for a minimum of ∼45 min to allow for de-esterification of the dye. A variable scan digital imaging system (Nikon NIS-Elements v4.3, Nikon GmbH) in connection with an upright microscope (Nikon Eclipsle FN-PT, Nikon GmbH) was employed for wide-field Na^+^ and pH imaging. SBFI was excited at 340 and 380 nm and emission collected >440 nm with a sampling rate of 0.5 Hz. BCECF was excited at 452 and 488 nm and emission collected between 490-575 nm (sampling rate of 0.2 Hz). Each choroid plexus was imaged for 2 min in HCO_3_^−^-aCSF to record the baseline. In Na^+^ imaging experiments, this was followed by wash-in of HCO_3_^−^-aCSF containing transporter inhibitors and imaging for five min in the presence of these (20 μM bumetanide ^29^, 2 mM ouabain ^30^, 30 μM S0859, or 10 μM cariporide ^31^, or corresponding DMSO vehicle). In pH imaging experiments, 5 mM NH_4_Cl was washed in for 10 min and imaging continued for another 18 min either in the absence or in the presence of the NHE1 inhibitor cariporide 10 μM). Analysis of fluorescence derived from SBFI or BCECF was performed from a region of interest (ROI) and background-correction carried out for each ROI and each wavelength. Afterwards, the fluorescence ratio was calculated (F_340_/F_380_ for SBFI and F_452_/F_488_ for BCECF) using the OriginPro Software (OriginLab Corporation v.9.0). The NKCC1 inhibitor bumetanide shows autofluorescence when excited at 340 nm, the “sodium-insensitive” wavelength of SBFI. Bath application of bumetanide thus resulted in a rapid, stepwise increase in fluorescence upon excitation at 340 nm in all investigated ROIs, including those chosen for background correction. Washout of bumetanide resulted in a likewise rapid decrease in fluorescence at 340 nm. Comparable fast changes in fluorescence emission following wash-in or wash-out of bumetanide were not observed upon excitation at 380 nm (“sodium-sensitive wavelength”), indicating negligible autofluorescence of bumetanide at this wavelength. In the following, autofluorescence was thus corrected for by subtracting the fluorescence recorded from a ROI in the field of view apparently free of SBFI-loaded cellular structures (“background”) from each individual ROI containing cellular structures, resulting in extinction of the fast autofluorescence-induced signal (see also ^32^). To determine changes in Na^+^ flux rates, linear regression analyses were performed on the rate of fluorescence change upon inhibitor infusion. Data analyses were carried out in a blinded fashion.

### ^86^Rb^+^ efflux

Acutely isolated choroid plexus from anesthetized rats and pigs was allowed to recover in 37 °C HCO_3_^−^-aCSF for 5-10 min (pig choroid plexus was dissected into smaller pieces) prior to exposure to the two isotopes in 37 °C HCO_3_^−^-aCSF for 10 min; ^86^Rb^+^ (1 µCi ml^−^, NEZ07200, Perkin Elmer) and ^3^H-mannitol (4 µCi ml^−1^, NET101, Perkin Elmer), the latter serves as an extracellular marker ^33^. The choroid plexus was briefly (15 s) washed in isotope-free HCO_3_^−^-aCSF prior to transfer into 1 ml HCO_3_^−^-aCSF containing vehicle, bumetanide (20 μM), or furosemide (1 mM). 0.2 ml of the surrounding aCSF was collected (and replaced) every 20 s and placed in scintillation vials. At the termination of the experiment, the choroid plexus was dissolved overnight at room temperature in 1 ml Solvable (6NE9100, Perkin Elmer) and combined with the remaining efflux solution. Determination of isotope content was obtained in Ultima Gold^TM^ XR scintillation liquid (6013119, Perkin Elmer) with a Tri-Carb 2900TR Liquid Scintillation Analyzer (Packard). For each time point (0, 20, 40, 60, 80 s), the ^86^Rb^+^ level was corrected for ^3^H-mannitol (extracellular background) and data are shown as the natural logarithm of the ^86^Rb^+^ level at time point, T, (A_T_) normalized to that of time point, 0, (A_0_) as a function of time. The ^86^Rb^+^ efflux rate constant (min^−1^) was given by the slope from linear regression analysis15,33.

### Immunohistochemistry

Anesthetized rats were perfusion fixed in 4% paraformaldehyde. Dissected ventricular regions (P21) or whole rat brains (9-week-old) were immersed in the fixative at 4 °C overnight. Whole brains were cryoprotected in 25 % sucrose and frozen in crushed solid CO_2_ and hippocampal brain regions were embedded in paraffin blocks. Cryostat sections (12-16 μm) were prepared and mounted on glass slides. To prepare the paraffin embedded regions, the glass slides were heated at 50-60 °C for 10 min to melt the paraffin. The heating was followed by boiling to retrieve antigens and expose the highest number of antigenic epitopes. Slides were immersed 2 x 5 min in xylene (534056, Sigma) and hydrated for 5 min in 99 % ethanol (100983, Merck), 5 min in 95 % ethanol, 5 min in 70 % ethanol, and 5 min in deionized water prior to incubation with citrate buffer (1.6 mM citric acid (251275, Sigma), 8.4 mM sodium citrate (1613859, Sigma) and boiled for 15 min. Cryostat sections were washed repeatedly in PBS and/or PBS-T (PBS with 0.03-0.25 % Triton-X100) for 5 min and blocked in 1-10 % serum diluted in PBS-T (blocking buffer) for 1-2 h at room temperature. This was followed by incubation for 30 min at room temperature and/or incubation overnight at 4 °C with primary antibodies (Rabbit anti-NKCC1: 59791, Abcam, 1:400 in blocking buffer; mouse anti-NKA α1: a6F AB 528092, DSHB, 1:300 in 1% serum in PBS-T). Sections were repeatedly washed for 5-10 min in PBS or PBS-T followed by incubation for 1-2 h with secondary antibodies (488-conjugated anti-mouse: 1:300 in serum-based (1 %) PBS-T or anti-rabbit: A-11034, LifeTech, 1:500 in blocking buffer). Negative controls with secondary antibodies were run in parallel. Subsequently, sections were washed by repeated 5 min intervals in PBS-T and/or PBS. Nuclei staining with Hoechst (1:10.000 in PBS) applied between initial washes or with ProLong Gold DAPI mounting medium (Dako). Micrographs were recorded using a Zeiss LSM710 point laser (Argon Lasos RMC781272) scanning confocal microscope with a Zeiss Plan-Apochromat 63×/numerical aperture (NA) 1.4 oil objective (Carl Zeiss, Oberkochen). All micrographs were sampled in a frame scan mode.

### Western blotting

Choroid plexus (lateral or 4^th^) was isolated in HEPES-aCSF, lysed in RIPA buffer (in mM: 150 NaCl, 50 Tris pH 8.0, 5 EDTA and 0.5% sodium deoxycholate, 0.1% SDS and 1% Triton X-100) with 0.4 mM pefabloc and 8 µM leupeptin, and sonicated (twice; 70 % power for 30 sec, Sonopuls, Bandelin) prior to protein concentration determination with the DC Protein Assay (Bio-Rad) according to the manufacturer’s instruction. 10-20 µg of total protein was loaded on precast SDS-PAGE gels (4-20% Criterion TGX, Bio-rad) and immobilon-FL membranes (Merck Milipore) employed for the transfer. Primary and secondary antibodies were diluted 1:1 in Odyssey blocking buffer (LI-COR): PBS-T. Primary antibodies: anti-GAPDH; AB2302 (Millipore, 1:5000), anti-NKCC1; S022D (MRC PPU Reagents, 2 µg ml^−1^), anti-KCC1 ^34^ (kind gift from Professor Thomas J. Jentsch, Max Delbrück Center For Molecular Medicine, Berlin, Germany, 1:500), anti-KCC2; 07-432 (Millipore, 1:1000), anti-KCC4 ^35^ (kind gift from Professor Jinwei Zhang, University of Dundee, UK, 2 μg ml^−1^), anti-NKA_1; a6F AB528092 (DSHB, 1:60), anti-NKA_2; 07-674 (Millipore, 1:500), anti-NKA_3; XVIF9-G10 (Thermo Fisher, 1:1000), anti-NHE1; sc-136239 (Santa Cruz, 1:500). Secondary antibodies, all 1:10,000; IRDye 680RD donkey anti-chicken (LI-COR, P/N 925-68075), IRDye 800CW goat anti-rabbit (LI-COR, P/N 926-32211), IRDye 800CW goat anti-mouse (LI-COR, P/N 926-32210), rabbit anti-sheep (Thermofisher, SA5-10060). Images were obtained by the Odyssey CLx imaging system and analyzed by Image Studio 5.2.5 (LI-COR).

### mRNA quantification

Total RNA was purified from rat choroid plexus embedded in RNAlater (R0901, Sigma) using the RNeasy micro kit and RNAase-free DNase set (both from Qiagen) according to manufacturer’s instructions. Reverse transcription was performed with 0.5 µ g cRNA using the Omniscript RT mini kit (Qiagen). cDNA amplication was carried out with Light-Cycler 480 SYBR Green I Master mix (Roche) on the Stratagene Mx3005P QPCR system (Agilent Technologies). Quantitative PCR settings: initial melting at 95 °C for 10 min, 40 amplification cycles at 95 °C for 10 s, primer annealing at 60 °C for 22 s, and elongation at 72 °C for 20 s. Amplification specificity was determined from melting curves and gel electrophoresis, see primer sequence in Supplementary Fig. 2. Primers were designed using NCBI’s pick primer software. The optimum concentration for each primer set was determined to 200□nM except for KCC4 (300 nM). Standard curves of 4× serial dilutions of plasmid cDNA or cDNA from reverse transcription of total RNA from either rat choroid plexus or rat total brain tissue were made in order to determine the amplification efficiencies for each of the utilized primer-sets, which were all between 90 and 110 %. Data analyses were performed in GenEx (MultiD Analyses AB). The genes of interest were normalized to the two reference genes ACTB and RPS18.

### RNA sequencing

Rat choroid plexus (lateral and 4^th^) was isolated in HEPES-aCSF and stored in RNAlater (R0901, Sigma) at −80 °C. The RNA extraction and library preparation were performed by Novogene Company Limited with NEB Next® Ultra™ RNA Library Prep Kit (NEB) prior to RNA sequencing (paired-end 150 bp, with 15Gb output) on an Illumina NovaSeq 6000 (Illumina). The 150 base paired-end reads were mapped to Reference genome Rnor_6.0 (*Rattus norvegicus*) using Spliced Transcripts Alignment to a Reference (STAR) RNA-seq aligner (v 2.7.2a) ^36^. The mapped alignment generated by STAR was normalized to transcripts per million (TPM) with RSEM (version 1.3.3) ^37,38^. Gene information was gathered with mygene (v 3.1.0) python library ^39,40^ (http://mygene.info), from which gene symbol, alias, and GO-terms ^41–43^ were extracted. The list of genes annotated as ‘transporters’ was obtained from the Guide to Pharmacology webpage (target and family list) ^44^ and employed to generate the list of membrane transport proteins. To exclude transport proteins in intracellular membranes, the generated list was filtered by initially removing the SLC25 mitochondrial solute carrier family, ATP5 family of mitochondrial membrane ATP synthase, and the ATP6V family of vacuolar H^+^-ATPase. Subsequently, all genes were included if no GO term were present or with associated GO terms; “integral component of plasma membrane” or “plasma membrane”, but only included genes annotated as “integral component of membrane”, “membrane” or “transmembrane”, which was not also annotated as “lysosome”, “endosome membrane”, “lysosomal”, “mitochondrion”, “mitochondrial”, “golgi apparatus”, “vacuolar”, or “endoplasmic”. STAR-RNA parameter settings for library build and mapping, together with all scripts for the gene annotation and analysis can be found at https://github.com/Sorennorge/MacAulayLab-RNAseq1.

### *Xenopus laevis* oocyte current recordings

cDNA encoding rat NBCe2 was purchased from GenScript and subcloned into the oocyte expression vector pXOOM. The construct was linearized downstream from the poly-A segment, and *in vitro* transcribed using T7 mMessage machine according to manufacturer’s instructions (Ambion). mRNA was extracted with MEGAclear (Ambion) and microinjected into defolliculated *Xenopus laevis* oocytes: 50 ng NBCe2 RNA/oocyte. *Xenopus laevis* oocytes were purchased from EcoCyte Bioscience and kept in Kulori medium (in mM): 90 NaCl, 1 KCl, 1 CaCl_2_, 1 MgCl_2_, 5 HEPES (pH 7.4) for 3-4 days at 19 °C prior to experiments. Oocyte two-electrode voltage clamp was performed at room temperature using the pCLAMP 10.4 software (Molecular Devices) together with the Clampator One amplifier (model CA-1B, Dagan Co.) and the A/D converter Digidata 1440A (Molecular Devices). Borosilicate glass capillary electrodes were pulled using a vertical pipette puller (PIP 5, Heka Elektronik) to a resistance of 1–3 MΩ when filled with 1 M KCl. Oocytes were subjected to a voltage clamp protocol, which comprised 200 ms steps from −120 mV to +40 mV in 20 mV increments from a holding potential of −60 mV. Currents were low-pass filtered at 500 Hz and sampled at 2 kHz. The bicarbonate-containing solution perfusing the oocyte chamber contained (in mM): NaCl 58.5; KCl 2.5; CaCl_2_ 1; MgCl_2_ 1; Na_2_HPO_4_ 1; NaHCO_3_ 24, HEPES 5 (pH 7.4), equilibrated with 5% CO_2_. The IC_50_ curve was made by perfusing the oocyte chamber with increasing concentrations of DIDS or S0859. Each concentration perfused the oocyte chamber for 4 minutes before the voltage clamp protocol was applied. Data analysis was performed in GraphPad Prism. Data were generated using at least two different oocyte batches. The current-voltage curves were obtained from the averaged currents at 160-200 ms of the voltage protocol and the IC_50_ curves obtained at +40 mV. The equation Y=Y_min_+(Y_max_−Y_min_)/(1+(IC_50_/X)^HillSlope) was used to fit the IC_50_ curve.

### Calculations of osmotic gradients

The osmotic gradient required to support the spontaneous transport of water across the choroidal epithelium (ΔOsm_req_, in Osm) can be calculated theoretically based on the functional water permeability across choroid plexus (*L_p_*, in cm s^−1^ Osm^−1^), the choroidal cross-sectional area (*A*, in cm^2^), and the CSF production rate (*V_p_*, in cm^3^ s^−1^):

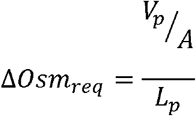

### Statistics

All data are shown as mean ± SEM. To test for statistically significant differences, we performed paired or unpaired two-tailed *t* tests, one or two-way ANOVAs followed by Dunnett’s or Tukey’s multiple comparisons *post hoc* test, as indicated in figure legends. P < 0.05 was considered statistically significant. All statistical analyses were carried out with GraphPad Prism (GraphPad Software).

### Data availability

We confirm that all data from this study will be available upon request.

## Results

### Choroid plexus water permeability

To quantify the CSF secretion as a function of the trans-epithelial osmotic gradient across the CSF-secreting choroid plexus, we determined the rate of CSF secretion in anesthetized rats during experimentally-inflicted modulation of the ventricular osmolality. The CSF secretion rate was determined with the ventriculo-cisternal perfusion technique, in which a dextran-based aCSF is infused in the right lateral ventricle of the rat brain, while collecting fluid from cisterna magna simultaneously^15^. Since the employed dextran, with its large molecular weight (Mw: 150,000), barely penetrates the ventricular walls at the employed time scale (Fig. 1a), the dextran dilution largely corresponds to newly formed CSF. As it is well established that the CSF secretion rate relies on a range of physiological parameters, such as blood pressure, heart rate, blood oxygenation, type of anesthesia etc., the experimental procedures were carried out with proper mechanical ventilation of the animal and monitoring of a range of physiological parameters, following determination of the optimal anesthesia regime and verification of stable blood electrolyte content and osmolality (see Supplementary Fig. 1 and Methods). The rate of CSF secretion in adult (9-week-old) male Sprague Dawley rats was obtained by infusing heated and equilibrated dextran-containing aCSF (with matching osmolality to that of rat CSF) into the lateral ventricle and recording the dextran content in the fluid collected from cisterna magna, see Fig. 1b for a representative experiment and inset for the summarized CSF secretion rate (6.8 ± 0.3 µl min^−1^, n = 28).

**Fig. 1.**
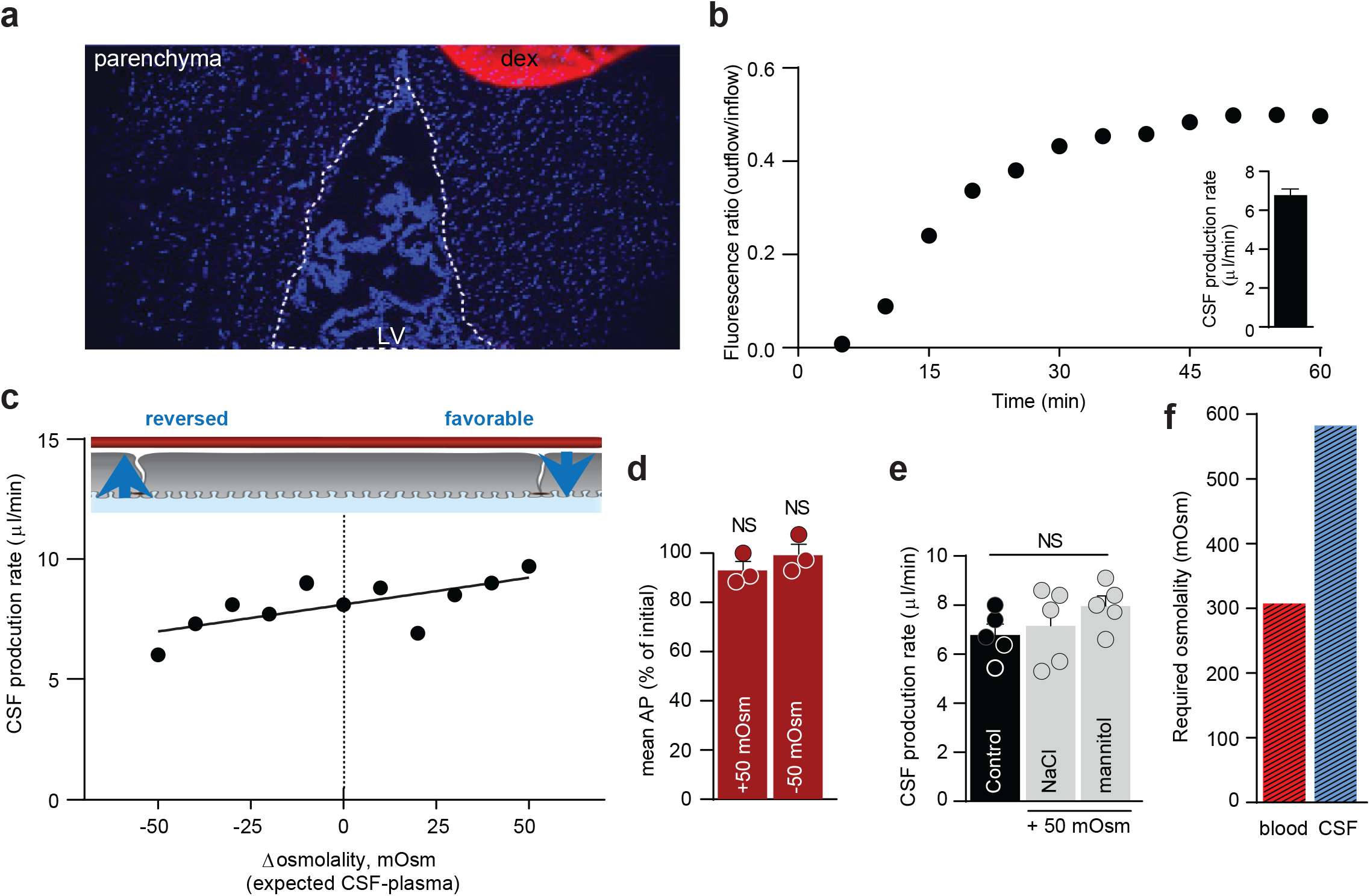
Negligible osmotic contribution to CSF secretion. **a**, Demonstration of dextran levels after an ended ventriculo-cisternal perfusion experiment in a rat. Representative sagittal cryostat sections of brains displaying the injected lateral ventricle (dotted line) and DAPI stained choroid plexus and parenchymal cells. A droplet of dextran with the concentration present within the ventricles during the experimental procedure was placed on the coverslip (red circle) and served as reference intensity. LV: lateral ventricle, dex: dextran. It is evident that the dextran had not diffused into the brain parenchyma to any detectable extent during the experimental procedure. **b,** Representative graph of the ratio of dextran (outflow/inflow) during a 1 h control ventriculo-cisternal perfusion performed in a rat. Inset denotes the average CSF production rate in control rats, n = 28. **c**, CSF production rate in rats subjected to varied trans-choroidal osmotic gradients by ventricular infusion of aCSF of osmolarities of ± 100 mOsm (compared to aCSF), expected to be diluted 1:1 with newly formed CSF. Each dot represents one rat, n = 11. The slope was determined from linear regression analysis (slope significantly different from 0, P < 0.05). **d,** Mean arterial pressure (AP) in rats subjected to ventriculo-cisternal perfusion upon ventricular infusion of aCSF with ± 100 mOsm (mannitol), providing an expected ventricular osmolarity of ± 50 mOsm. Data are presented as the AP (in %) at the end of the experiment compared to the initial AP measurement, n = 3, paired t-test. **e,** CSF secretion rates obtained from ventriculo-cisternal perfusion of rats upon elevated (∼50 mOsm) ventricular osmolarity with either mannitol or NaCl as the osmotic agent, n = 5. One-way ANOVA followed by Tukey’s multiple comparisons test, NS = not significant. **f,** Required osmotic gradient to sustain a CSF production rate of 6.8 μl min-1 given the osmotic water permeability of the choroid plexus: Osmolality of CSF should exceed that of blood with ∼280 mOsm.

To determine the CSF secretion rate as a function of the osmotic gradient across the choroid plexus, CSF secretion was measured with varying osmolality of the infused aCSF (up to ± Δ100, mOsm, of that of the plasma osmolality with a predicted 1:1 dilution with the native CSF; ± Δ50 mOsm). The rate of CSF secretion increased slightly in response to increasing ventricular osmolality (Fig. 1c). Each mOsm elevation in the ventricular aCSF led to a 0.4 % increase in CSF secretion rate (0.023 ± 0.008 µl × min^−1^× mOsm^−1^, P < 0.05) with no effect of altered ventricular osmolality on the mean arterial blood pressure (in % of the value at the initiation of the experiment: +50 mOsm = 93 ± 4 % (P = 0.19), −50 mOsm = 99 ± 4 % (P = 0.84), n = 3 of each, Fig. 1d). To retain the ionic composition of the aCSF in this series of experiments, the aCSF osmolality was adjusted with addition or removal of mannitol from the isosmotic control solution (as an inert osmolyte, see Methods). The CSF secretion rate was similar whether the hyperosmolar challenge arose from addition of mannitol (8.0 ± 0.4 µl min^−1^) or NaCl (7.2 ± 0.7 µl min^−1^, n = 5 of each P = 0.55, Fig. 1e).

To obtain the passive osmotic water permeability of the rat choroid plexus epithelium, we assessed the cross-sectional surface area of the choroid plexuses (lateral + 4^th^) based on their wet-weight. 88% of the choroidal weight has been estimated to consist of epithelium ^45^, which gives 4.6 ± 0.4 mg choroid plexus epithelium per rat (n = 5). The choroidal volume thus approximates 4.6 × 10^−9^ cm^3^, assuming a cell density of 1 g/cm^3 46^. Given a cell height of ∼10 μm ^47^, the cross sectional area of (lateral and 4^th^) choroid plexus was 4.6 cm_2_, which aligns with ^45^ upon age correction of the rats ^48^. Taken together with the CSF secretion rate as a function of the osmotic gradient (0.023 ± 0.008 µl × min^−1^ × mOsm^−1^, Fig. 1c), the passive, trans-epithelial water permeability of the rat choroid plexus yields an L_p_ = 8.7 × 10^−5^ cm s^−1^ Osm^−1^. If the CSF should be produced entirely from conventional osmotic driving forces with the given osmotic water permeability, the theoretical ventricular osmolality required to drive the basal CSF secretion rate of 6.8 µl min^−1^ can be calculated to ∼280 mOsm higher than that of the plasma (= ∼ 590 mOsm, Fig. 1f)

### Absent trans-choroidal osmotic gradient

Based on the theoretical requirement for a large ventricular osmolality to drive CSF production of the given rate by conventional osmotic water flux, we determined the trans-epithelial osmotic gradient across the rat choroid plexus epithelium. To limit confounding alterations in electrolyte content (and thus osmolality), the experimental procedures were designed to limit the time from anesthesia induction to CSF/blood sample collection. With the swift protocol, we obtained osmolalities of 306 ± 4 mOsm for plasma and 307 ± 2 mOsm for CSF (sampled from cisterna magna; n = 6, Fig. 2a). The osmolality of these fluids surrounding the choroid plexus is thus equivalent (P = 0.78), illustrating that there is no trans-choroidal osmotic gradient to drive CSF secretion in rats. To verify this observation in a larger animal, we determined the osmolality of ventricular CSF (collected from cisterna magna) and blood extracted from anesthetized and mechanically ventilated pigs. The osmolality of pig blood (290 ± 1 mOsm) and CSF (291 ± 2 mOsm) was similar (n = 7, P = 0.22, Fig. 2b), as was the case for serum (293 ± 2 mOsm) and lumbar CSF (296 ± 2 mOsm) collected from healthy human individuals screened, but subsequently cleared for, normal pressure hydrocephalus (n = 14, P = 0.09, Fig. 2c). Note that regular anti-coagulant (EDTA)-coated blood sample tubes produce an elevation of the blood osmolality (Supplementary Fig. 3). All blood samples were therefore collected either in the absence of anticoagulants (human) or in the presence of heparin (rat and pig), which did not significantly alter the plasma osmolality (Supplementary Fig. 3a-b). In summary, the osmolality of the bulk CSF is not elevated above that of the blood in either of the tested species. The osmotic gradient across the choroid plexus epithelium *in vivo* is thus orders of magnitude too small to drive CSF secretion by conventional osmosis. CSF secretion against osmotic gradients

**Fig. 2.**
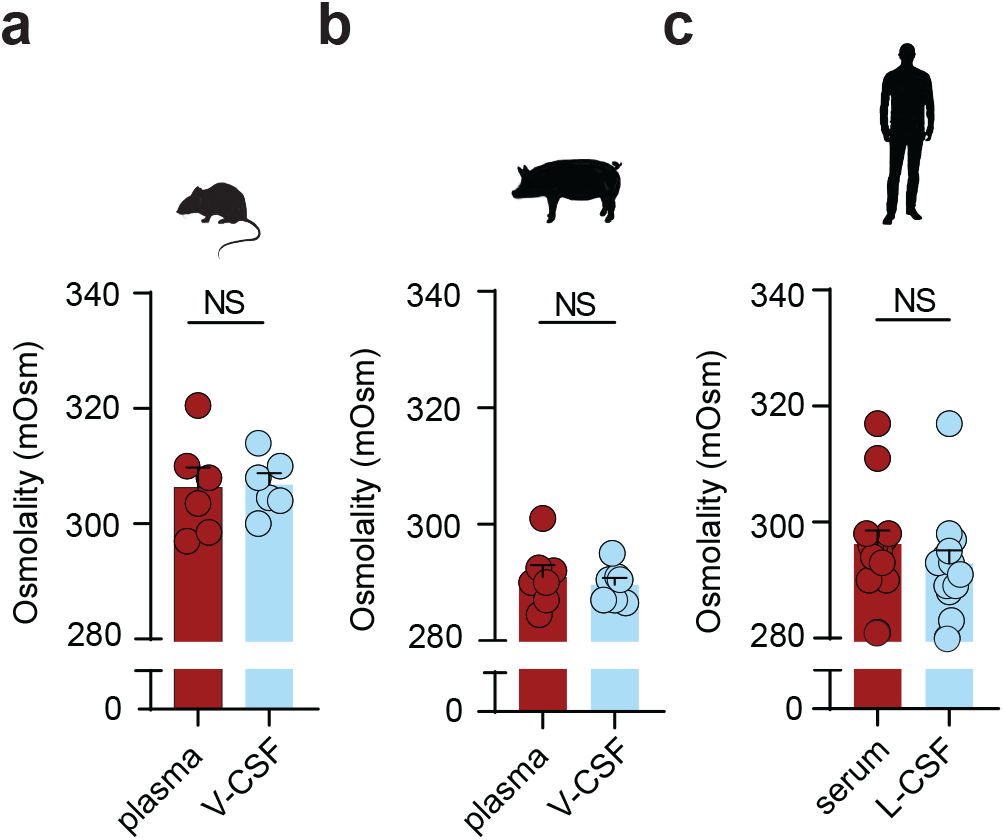
Osmolality of blood and cerebrospinal fluid. Osmolality of blood (plasma or serum) and ventricular (V) or lumbar (L) cerebrospinal fluid from rats, n = 5 (**a**), pigs, n = 7 (**b**) and humans, n = 14 (**c**). Paired two-tailed *t*-test, NS = not significant.

It is evident from Fig. 1c that increased ventricular osmolality slightly elevates the rate of CSF secretion. However, the left side of the graph depicts an experimental scenario in which the plasma osmolality is higher than that of the CSF. Under these conditions, the osmotic gradient would favor fluid flow *from the ventricles to the vasculature*. Nevertheless, CSF secretion continues, even when the choroid plexus faces an *oppositely directed* osmotic gradient of approximately 50 mOsm, a condition under which the CSF secretion is reduced a mere 20%. This finding illustrates that CSF can be secreted independently of, and even against, a sizeable bulk osmotic gradient. Taken together with the lack of a detectable osmotic gradient across the choroid plexus epithelium and its low trans-epithelial osmotic water permeability, we conclude that the CSF secretion must occur by mechanisms other than conventional osmotic water transport.

### Local osmotic gradients cannot explain CSF secretion

The luminal surface membrane of the cuboidal-to-columnar choroid plexus epithelium is covered by abundant microvilli, which serve to increase the luminal surface area. The inter-microvillar space might serve as a protected area where solutes transported across the epithelium accumulate and create a local osmotic gradient driving fluid across the tissue even in the absence of an osmotic gradient in the bulk solutions surrounding the epithelial cell membrane. To determine if such a mechanism could account for the ability of the choroid plexus to secrete CSF in the absence of a bulk osmotic gradient, we created a mathematical model of solute-linked water transport in the inter-microvillar space of the luminal membrane of the choroid plexus.

To obtain the anatomical parameters of choroid plexus from rats of the species and age employed for the physiological experiments, we conducted morphometric analysis following transmission and scanning electron microscopy of the acutely extracted tissue. Scanning electron microscopy reveals the gross structure of part of the choroid plexus (Fig. 3a); at higher magnification, individual tissue fronds are seen to be covered by an irregular epithelium with abundant microvilli on the luminal surface (length 1.71 ± 0.09 μm, n = 45 from ten different cells, Fig. 3b-c). The microvilli are typically cylindrical at the base (radius 0.06 ± 0.01 μm, n = 13) but increase in diameter towards the tip to end in a small bulbus (radius 0.12 ± 0.01 μm, n = 8; see arrows in Figure 3b-d). The extent of the brush border is somewhat smaller (ca. 1.5 μm) than the length of microvilli because of their flexibility and interdigitation (Fig. 3b-c). The spatial density of microvilli was estimated from the linear density on TEM images (Fig. 3e) to be 18 microvilli per µm^2^, which altogether increased the surface area of the luminal membrane 12-fold. With the 4.6 cm^2^ apparent luminal surface obtained above, the estimated true membrane area arrives at 57 cm^2^ in the 9-week-old Sprague Dawley rat.

**Fig. 3.**
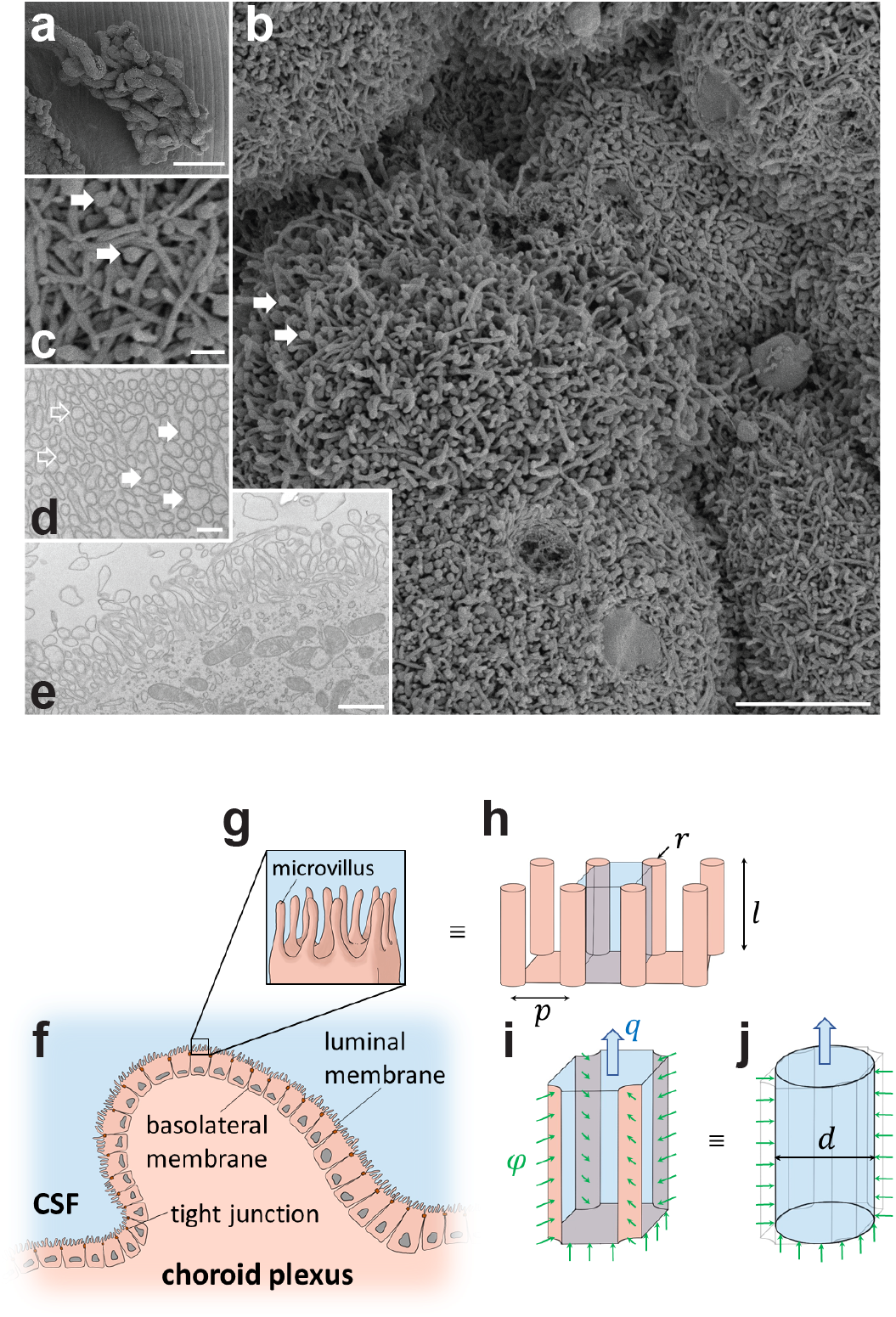
Ultrastructure of the choroid plexus luminal membrane and a schematic mathematical model domain derivation. **a-c**, Scanning electron microcopy images of rat choroid plexus. **a**, the typical frond-like arrangement of the epithelium dictated by the ensheathment of blood vessels in the underlying connective tissue. **b**, At higher magnification, the abundant microvilli and the loose brush border that they form is visible. Arrows point to the terminal bulbous tip of the microvilli, also shown at higher magnification in **c**, which also reveals the strong interdigitation of the microvilli. **d-e**, Transmission EM images of the epithelium show the space filling of microvilli cut perpendicularly a distance from the luminal membrane (**d**) and the extent of the brush border (**e**). Note change in diameter of microvilli from base (open arrows) to tip (filled arrows). Bars, a; 250nm, b; 5 µm, c; 500 nm, d; 250 nm, and e; 1 µm. **f**, Local cross-section of the choroid plexus. Luminal and basolateral membranes, as well as tight junctions are indicated. **g**, Magnified view of the luminal epithelial cell surface showing microvilli and the inter-microvillar space, where a standing osmotic gradient might exist. **h**, Geometric approximation of the luminal cell surface, with microvilli represented as cylinders with radius *r* and length *l*, and separated by a distance of *p*. The blue region shows one inter-microvillar space. **i**, A single inter-microvillar space with interfaces to the four adjacent microvilli and to the base of the luminal membrane of the epithelial cell. The green arrows indicate solute flux, φ through these interfaces. The solute flux is produced by transport mechanisms on the luminal membrane. Potential accumulation of solutes in the inter-microvillar space might yield osmotic forces that draw fluid from the choroid plexus epithelium, resulting in a CSF secretion rate of *q* per such space. The overall CSF production rate by this mechanism is obtained by multiplying *q* with the number of inter-microvillar spaces on the entire choroid plexus surface. **j**, Circular cylindrical channel with diameter *d* that is hydrodynamically equivalent to the inter-microvillar space shown in (i). A one-dimensional standing osmotic gradient model was derived from this geometry.

For the model, we used the simplified anatomic representation shown in Fig. 3f-j. We assumed that all solutes in the ventricular CSF enter through the inter-microvillar space, likely overestimating the local osmotic gradient, and that the solutes are removed by bulk CSF drainage at the same rate as they enter. Given an isotonic CSF secretion rate of 6.8 µl min^−1^, a bulk CSF osmolality of 307 mOsm (Fig. 2a), and a CSF density of 1.00 g ml^−1 49^, the ion transfer rate across the entire luminal membrane amounts to 3.48 ×10^−5^ mmol s^−1^ (see Supplementary Data). Our model only accounts for the luminal membrane, but the trans-choroidal osmotic water permeability obtained above includes the effect of the two epithelial membranes. Each membrane’s osmotic water permeability is double the measured value, since the membranes are configured in series. Accordingly, the osmotic water permeability of the luminal membrane (true area) was calculated to be 1.4 × 10^−5^ cm s^−1^ Osm^−1^. With these values, our model predicts that inter-microvillar local osmotic forces may account for less than 0.1% of the observed CSF secretion rate (0.004 µl/min of the measured 6.8 µl min^−1^). Our mathematical model thus suggests that CSF secretion does not originate from local osmotic gradients built up between the choroidal microvilli.

### Choroidal transport proteins in the luminal membrane

RNAseq on the rat choroid plexus identified highly expressed electrolyte transporters in choroid plexus that could contribute to ion and fluid secretion across the luminal membrane: the Na^+^/K^+^-ATPase (α1), NKCC1, and the bicarbonate transporter NBCe2 all appeared in the top 6% highest expressed transcripts encoding plasma membrane ion transporters: The Na^+^/K^+^-ATPase subunit 1 ranked as number 4 (TPM = 516; represents ∼4% of all plasma membrane transporters in choroid plexus), NBCe2 ranked as number 5 (TPM = 504; represents ∼4% of all transporters in choroid plexus), and NKCC1 as number 13 (TPM = 230, represents ∼2% of all transporters in choroid plexus) out of a total of 233 transcripts encoding choroidal plasma membrane ion transporters (Fig. 4a). The Na^+^/H^+^ exchanger NHE1 appeared further down the rank list (number 131 out of 233 transcripts, with a TPM = 7; represents ∼0.05% of all transporters in choroid plexus). These findings were essentially verified with qPCR, although with a slightly lower, but still robust, expression of NBCe2 in the lateral choroid plexus (Fig. 4b) and in the 4^th^ choroid plexus (Supplementary Fig. 4a).

**Fig. 4.**
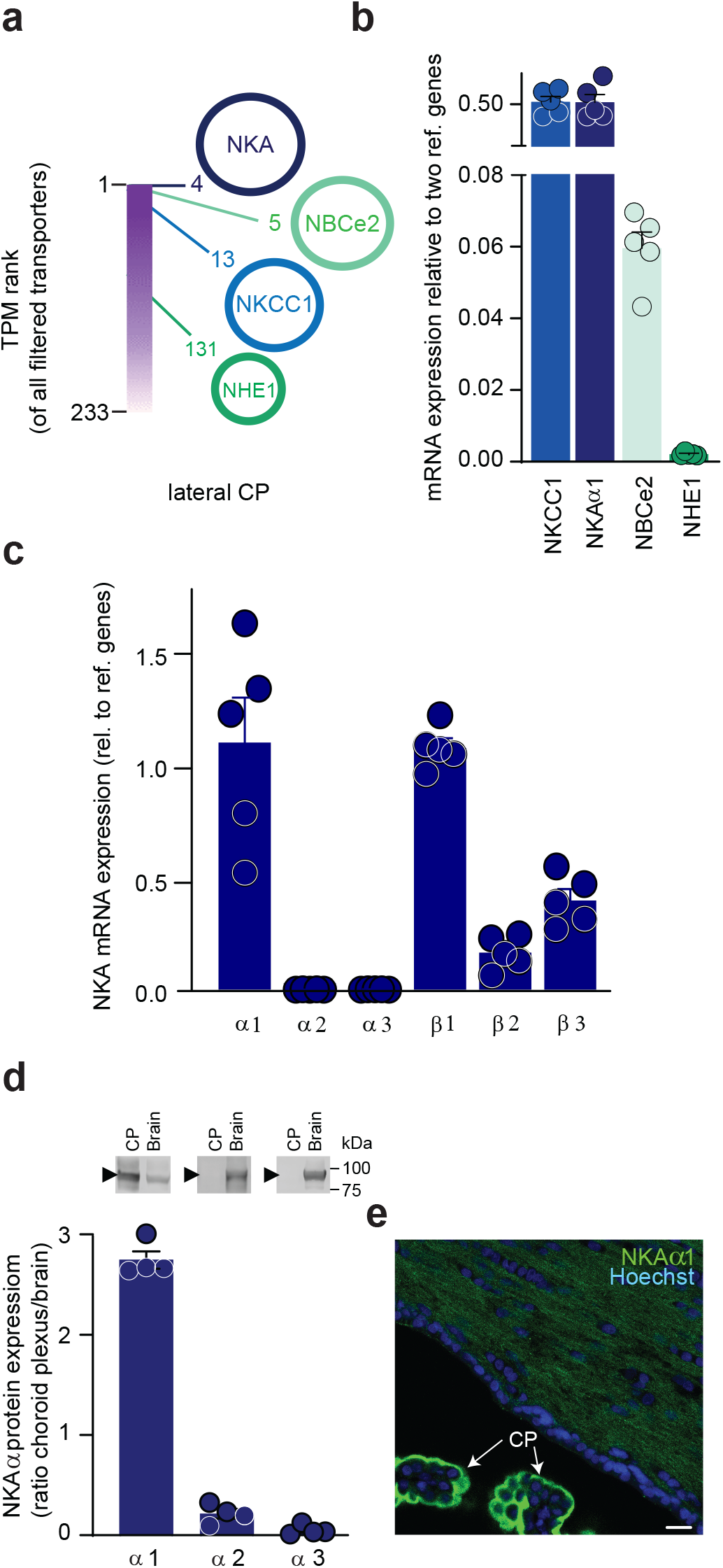
Quantitative mapping and expression of choroid plexus transporters. **a**, RNAseq of rat choroid plexus revealed transport proteins with high transcript abundance. Of a total of 233 genes in the filtered ion transporter category, see methods, the luminally expressed Na^+^/K^+^ ATPase (NKA) subunit u:1, NBCe2, and NKCC1 were in the top 6% of highest expressed transporters with rank numbers of 4, 5, and 13, with NHE1 at a much lower abundance; ranked 131. **b**, Relative mRNA expression levels of NKCC1 (0.52 ± 0.06), the NKA subunit α1 (0.52 ± 0.08), NBCe2 (0.06 ± 0.01), and NHE1 (0.002 ± 0.000) in rat choroid plexus, n = 5. **c,** mRNA expression levels in rat lateral of Na^+^/K^+^-ATPase subunit *α*1, *α*2, *α*3, *β*1, *β*2, and *β*3 relative to reference genes, n = 5. **d**, Quantification of Na^+^/K^+^-ATPase subunit α1, α2, and α3 in choroid plexus and brain normalized to GAPDH loading control. Data are shown as choroid plexus expression relative to brain expression, n = 4. Inset: Representative Western blots of Na^+^/K^+^-ATPase subunit α1, α2, and α3 in lateral choroid plexus or whole brain tissue (see intact Western blots in Supplementary Fig. 4d). **e**, Immunostaining of Na^+^/K^+^-ATPase subunit α1 (green) in rat choroid plexus (CP) and brain parenchyma. Cell nuclei visualized with Hoechst (blue), scalebar = 20 μm.

The NKCC1, NBCe2, NHE1, and Na^+^/K^+^-ATPase (α1) localize to the luminal choroidal membrane in all tested species ^50–54^, here verified for NKCC1 in the rat choroid plexus (Supplementary Fig. 4b). The α1 subunit of the Na^+^/K^+^-ATPase is the dominant catalytic subunit in choroid plexus, with the α2 and α3 transcripts appearing at much lower levels (Fig. 4c and Supplementary Fig. 4c). The high β1 transcript level suggests this accessory subunit as the dominant, although both β3 and, to a lesser extent, β2 are expressed in the rat choroid plexus (Fig. 4c and Supplementary Fig. 4c) and may serve to modulate the pump function ^55^. Notably, the abundance of Na^+^/K^+^-ATPase α1 protein is 3-fold higher in choroid plexus than in brain parenchyma, whereas the glial α2 and the neuronal α3 subunits ^55^ are practically absent in choroid plexus (Fig. 4d-e, and Supplementary Fig. 4d).

### Contribution of transporters to CSF secretion

Quantification of the individual contribution to CSF secretion of these luminally expressed transport mechanisms was obtained with a pharmacological strategy (Fig. 5a). Efficient and well-established inhibitors exist for NKCC1 (bumetanide), the Na^+^/K^+^ ATPase (ouabain), and NHE1 (cariporide), but the bicarbonate transporters are notoriously difficult to inhibit specifically and drug potencies not well defined. Prior to initiating the *ex vivo* and *in vivo* experimentation, we therefore determined the inhibitor concentration profile of the NBC inhibitors DIDS ^56^ and S0859 ^57^, the latter employed in LI-COR experiments and [Na^+^]_i_ determinations due to DIDS’ autofluoresence in these spectra. To this end, rat NBCe2 was expressed heterologously in *Xenopus laevis* oocytes and its activity monitored by two-electrode voltage clamp. NBCe2-mediated membrane currents were abolished with 30 µM S0859 (IC_50_ of 10.7 ± 0.9 µM, n = 9, Supplementary Fig. 5a-c) and with 300 µM DIDS (IC_50_ = 4.5 ± 1.8 µM, n = 6, Supplementary Fig. 5d-f).

**Fig. 5.**
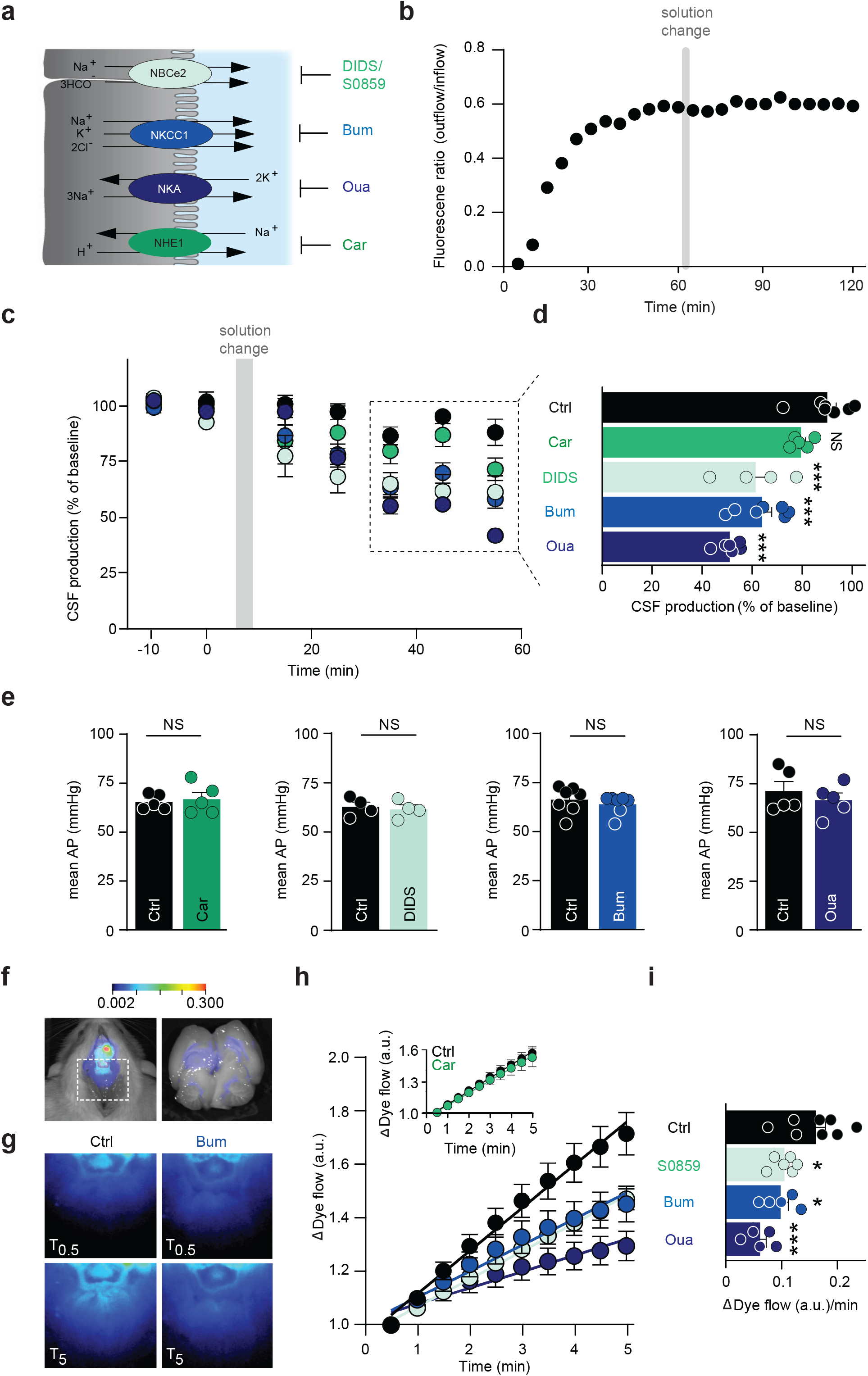
Three transporters as key contributors of CSF secretion. **a**, Illustration of the luminal ion transport proteins: NBCe2, NKCC1, NKA, and NHE1 in choroid plexus (grey) and their inhibitors DIDS/S0859, bumetanide, ouabain, and cariporide. CSF is shown in blue. **b,** Representative time control experiment with ventriculo-cisternal perfusion in rats depicting the ratio of dextran (outflow/inflow) as a function of time with a solution change to an identical solution. **c,** Quantification of the CSF production rate by ventriculo-cisternal perfusion in rats treated with intraventricularly-delivered inhibitors of NKCC1 (bumetanide, n = 7), NKA (ouabain, n = 6), NHE1 (cariporide, n = 5), NBCe2 (DIDS, n = 4) or control solution (n = 7) at solution change (grey box). **d,** Summarized data obtained after 35-55 min of inhibitor/vehicle exposure, with statistical significance determined by one-way ANOVA, followed by Dunnett’s multiple comparisons test. Asterisks denote difference from control condition (vehicle), ***; P < 0.001, NS = not significant. **e,** Mean arterial blood pressure (AP) obtained during the ventriculo-cisternal perfusion experiments is illustrated before and after (55 min) intraventricular exposure of the pharmacological inhibitors. Statistical significance was determined with paired *t*-test. NS = not significant. **f,** Illustration of IRDye 800 CW carboxylate dye injected in the right lateral ventricle shown in pseudo-colors superimposed on a white light image of the rat. The dye intensity is quantified as a function of time in the white box placed at lambda (left panel). Right panel illustrates a superimposed image of ventricular fluorescence on a white light image of a rat brain cut in the midsagittal plane after ended live imaging. The intensity scale is in arbitrary units. **g**, Representative images of the dye content at T0.5 min and T5 min after placement in fluorescent scanner in control rats (Ctrl) or bumetanide-treated rats (Bum). **h**, Live imaging of intraventricular IRDye 800 CW carboxylate dye following treatment with bumetanide (NKCC1 inhibition, n = 5), ouabain (NKA inhibition, n = 5), or S0859 (NBCe2 inhibition, n = 6) or control (vehicle) solution (n = 8). Inset: control: 0.12 ± 0.01 a.u./min,n = 7, vs. cariporide (NHE1 inhibition): 0.12 ± 0.02 a.u./min, n = 6. **i,** Quantification of the dye flow rate determined from linear regression analysis of **h** over the 5 min with statistical significance determined by one-way ANOVA followed by Dunnett’s multiple comparisons test. Asterisks denote difference from control condition, ***; P < 0.001, *; P < 0.05.

To determine the quantitative contribution of the luminally-expressed transport proteins, the ventricular-cisternal perfusion method was employed on anesthetized rats with inclusion of the various inhibitors. To obtain high level of fidelity, each rat served as its own control. Fig. 5b illustrates a time control experiment, in which the infused aCSF was replaced with an identical solution when the dextran dilution had stabilized (1 h) and the CSF secretion rate thus reliably monitored. The CSF secretion rate remained stable over the 2 h experimental window (Fig. 5b-c). With an expected ventricular dilution of at least 1:1 of the intraventricularly infused inhibitor concentrations in the ventriculo-cisternal perfusion, the inhibitor concentrations of the infused aCSF were adjusted to increase the probability of targeting the candidate transport mechanisms in both lateral ventricles. We thus expect maximal ventricular drug concentrations of 50 µM bumetanide, 2.5 mM ouabain, 50 µM cariporide, and 750 µM DIDS, but with lower concentrations near the (contralateral) choroidal surfaces. Each rat served as its own control with introduction of inhibitor (or vehicle) at a point of stable baseline. The CSF secretion was reduced by 36 ± 4 % with NKCC1 inhibition (bumetanide; n = 7, P < 0.001), 49 ± 2 % with Na^+^/K^+^-ATPase inhibition (ouabain; n = 6, P < 0.001), 28 ± 4 % with NBCe2 inhibition (DIDS; n = 4, P < 0.001), but unchanged with NHE1 inhibition (cariporide; n = 5, P = 1.0, Fig. 5c-d).

Several of these transporters may be directly or indirectly implicated in setting the blood pressure, alteration of which could indirectly affect CSF secretion. The mean arterial blood pressure was therefore recorded in parallel with the ventriculo-cisternal perfusion. The blood pressure (prior to inhibitor infusion) was 67 ± 1 mmHg, n = 31. After infusion of the inhibitor, the blood pressure was (in % of own control) 97 ± 2 for bumetanide (n = 7, P = 0.13), 94 ± 4 for ouabain (n = 5, P = 0.16), 98 ± 2 for DIDS (n = 4, P = 0.31), and 102 ± 3 for cariporide (n = 5, P = 0.47) (Fig. 5e) and, thus, remained stable during the procedure.

Although inhibition of these ubiquitously expressed transport mechanisms did not alter the mean arterial blood pressure, they, upon longer experimental regimes, could penetrate the brain parenchyma and indirectly affect CSF secretion in unpredictable manners. For swifter recordings of the CSF dynamics with a less-invasive approach, we took advantage of the LI-COR Pearl Trilogy small animal imaging system ^15^. This procedure allows for visualization of the two-dimensional rostral to caudal movement of fluorescent dye injected into the ventricles of the anesthetized rats within minutes of injection (Fig. 5f, left panel, for verification of the dye reaching the ventricular system, see Fig. 5f, right panel). The initial movement of the dye from the lateral ventricle into the region of interest (Fig. 5f-g) was quantified by linear regression (Fig. 5h) and employed as a proxy for CSF production (although a component is attributed to dye diffusion). After inclusion of the various inhibitors, the rate of control dye movement (0.16 ± 0.18 a.u. min^−1^, n = 8) was reduced by 39 % with NKCC1 inhibition (bumetanide: 0.10 ± 0.01 a.u. min^−1^, n = 5), 62 % with Na^+^/K^+^-ATPase inhibition (ouabain: 0.06 ± 0.01 a.u. min^−1^, n = 5), and 35 % with NBCe2 inhibition (S0859: 0.11 ± 0.01 a.u. min^−1^, n = 6) (Fig. 5h-i). Inhibition of NHE1 had no effect on the CSF flow (control: n = 7, cariporide: n = 6, P = 0.75, Fig 5h inset). These results qualitatively mimic those obtained with the ventriculo-cisternal perfusion method and indicate that the Na^+^/K^+^-ATPase, NKCC1, and NBCe2 transporters contribute significantly to CSF secretion. Taken together, these data suggest that inhibition of individual choroidal transporters, the Na^+^/K^+^-ATPase, NKCC1, and NBCe2, affects the rate of CSF secretion in a direct manner, while NHE1 did not contribute to this fluid secretion.

### Quantification of choroidal Na^+^ efflux

To rule out any indirect effect of these pharmacological inhibitors, we took advantage of the CSF secretion being correlated with the net flux of Na^+^ across choroid plexus ^58^. To fashion an experimental design with which to quantify each candidate transporter’s contribution to the choroidal net efflux of Na^+^, as a proxy of CSF secretion, we employed acutely excised choroid plexus tissue loaded with the Na^+^-sensitive fluorescent dye SBFI (Fig. 6a). The [Na^+^]_i_ dynamics were monitored by wide-field SBFI imaging upon acute application of the transport inhibitors. The SBFI dynamics demonstrated an immediate *increase* in [Na^+^]_i_ upon inhibition of NKCC1 (20 µM bumetanide: 0.034 ± 0.005 a.u. min^−1^, n = 7, P < 0.01), the Na^+^/K^+^-ATPase (2 mM ouabain: 0.022 ± 0.004 a.u. min^−1^, n = 6, P < 0.01), or NBCe2 (30 μM S0859: 0.023 ± 0.006 a.u. min^−1^, n = 5, P < 0.05), while the [Na+]i remained stable upon NHE1 inhibition (10 μM cariporide: 0.002 ± 0.001 a.u. min^−1^, n = 6, P = 0.07, Fig. 6b-c). This inhibitor-mediated [Na+]i elevation in choroid plexus epithelial cells demonstrates that NKCC1, the Na^+^/K^+^-ATPase, and NBCe2 work in the direction promoting Na+ efflux across the luminal membrane of choroid plexus. Verification of NHE1 protein expression in rat choroid plexus was demonstrated with Western blotting (Fig. 6d). The functional luminal membrane localization of NHE1 was visualized with fluorescence-based pH monitoring of choroid plexus challenged with an acid load ^59^, the return of which was delayed in presence of the NHE1 inhibitor cariporide (Fig. 6e-f). Taken together, the complementary *in vivo* and *ex vivo* data illustrate key roles for NKCC1, NBCe2, and the Na^+^/K^+^- ATPase in CSF secretion across the luminal membrane with no observed contribution from NHE1 in CSF secretion.

**Fig. 6.**
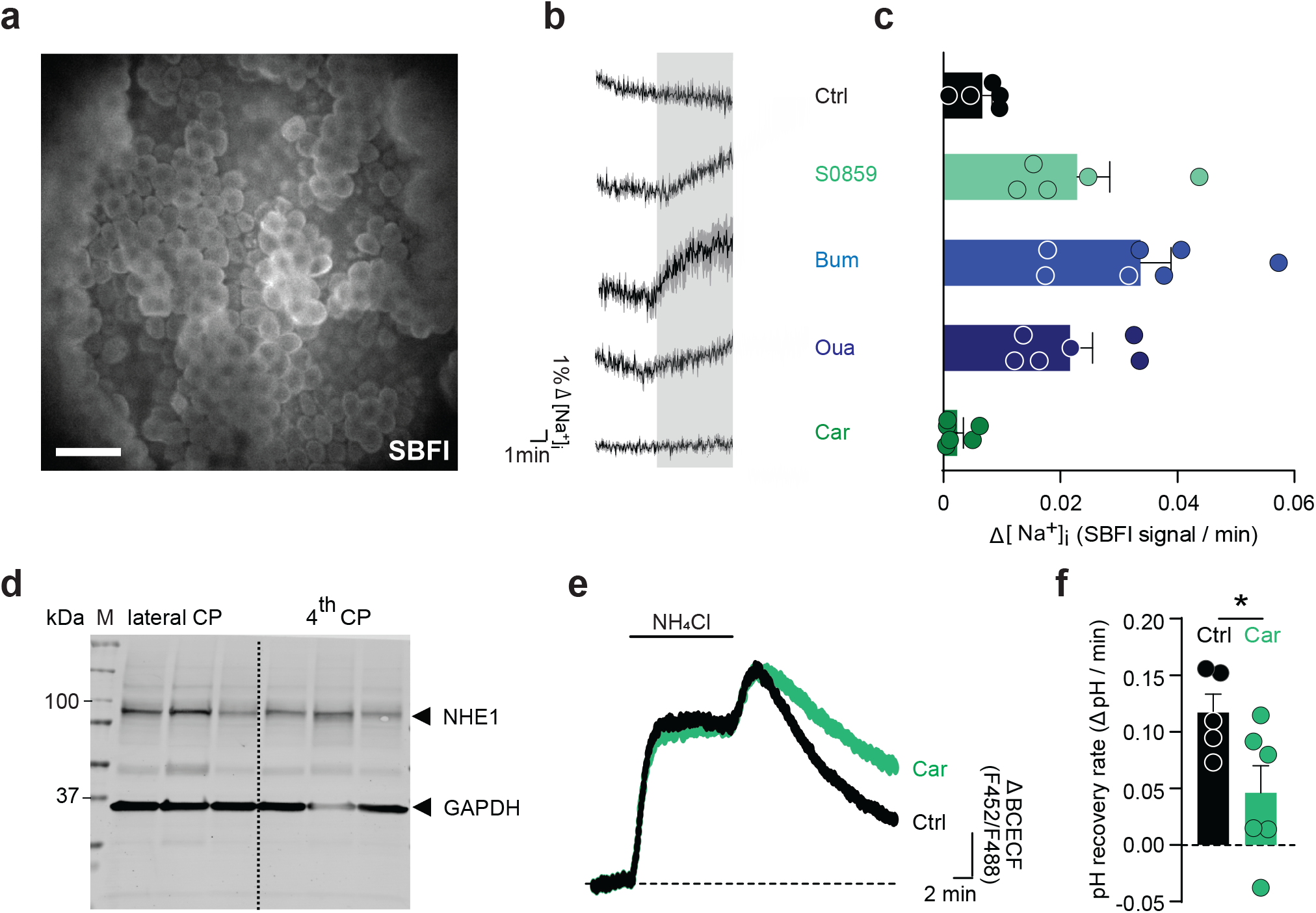
Transporter-mediated changes in choroidal [Na^+^]_i_ and pH. **a,** Representative SBFI fluorescence image of the lateral choroid plexus taken with a wide-field fluorescence microscope. Scale bar = 40 µm. **b,** Changes in SBFI fluorescence ratio, reporting changes in the intracellular [Na^+^] in choroid plexus in control solution (n = 5) or upon treatment with S0859 (NBCe2 inhibition, n = 5), bumetanide (NKCC1 inhibition, n = 7), ouabain (NKA inhibition, n = 6), or cariporide (NHE1 inhibition, n = 6) for 5 min. **c,** Quantification of the change of the SBFI signal (in **b**) as a function of time by linear regression, n = 5-7. **d,** Representative Western blot illustrating the protein expression of NHE1 in rat choroid plexus from lateral and 4^th^ ventricles. GAPDH was included as a loading control. **e,** Peak-normalized changes in BCECF ratio (arbitrary units) upon application NH_4_Cl, illustrating differences in the rate of recovery from acidification after removal of NH_4_Cl. Choroid plexus was perfused with aCSF containing 5 mM NH_4_Cl (indicated with black line) in the presence or absence of cariporide (NHE1 inhibitor). **f,** pH recovery rates determined from linear regression analysis on the period right after the maximal acid load in the presence (n = 6) and absence of cariporide (n = 5). Statistical significance determined with Student’s t-test. *P < 0.05.

### Outward-directed NKCC1, not KCC, transport

NKCC1’s ability to contribute directly to CSF secretion via its ability to transport water hinges on its transport being outwardly-directed. To resolve the NKCC1 transport direction, we determined the choroidal ^86^Rb^+^ efflux rate constant in acutely isolated, ^86^Rb^+^-loaded, rat choroid plexus. ^86^Rb^+^ can replace K^+^ at its binding site and thus act as a tracer for the K^+^ transport. The ^86^Rb^+^ efflux rate constant from rat choroid plexus was diminished by ∼60 % in the presence of bumetanide (0.40 ± 0.02 min^−1^ in control, n = 7, vs. 0.15 ± 0.01 min^−1^ in bumetanide, P < 0.001, n = 5, Supplementary Fig. 6a), demonstrating the bumetanide-sensitive outward flux of ^86^Rb^+^. NKCC1-mediated K^+^ efflux was verified in acutely isolated pig choroid plexus, in which inclusion of bumetanide led to a 68% decrease of ^86^Rb^+^ (K^+^) release (control: n = 6, bumetanide: n = 7, P < 0.001, Supplementary Fig. 6b). Different isoforms of the related K^+^/Cl^+^ cotransporters have been proposed to be expressed in the choroid plexus and potentially contribute to CSF secretion ^14,60,61^. Semi-quantification of their transcripts in rat choroid plexus by qPCR demonstrated expression of KCC1, albeit at lower levels than that of NKCC1, whereas the transcript levels of KCC2-4 were near/below detection range (Supplementary Fig. 6c). The expression of NKCC1 and KCC1, and absence of other KCCs, was verified at the protein level in rat and pig choroid plexus by Western blotting (Supplementary Fig. 6c, inset and 6d). To determine whether functional KCC expression occurred in the luminal membrane of choroid plexus, we included furosemide, a common blocker of NKCC1 and KCCs, in the ^86^Rb^+^ efflux assays. The inhibitory potential of furosemide did not exceed that of bumetanide in choroid plexus from both rat and pigs (Supplementary Fig. 6a-b). Taken together, these findings identify NKCC1 as an outwardly-directed transport mechanism in acutely isolated choroid plexus from rats and pigs, and demonstrate a lack of KCC activity at the luminal membrane in this tissue.

### NKCC1-mediated transport of water

To demonstrate NKCC1-mediated movement of water in rat choroid plexus, isolated rat choroid plexus was loaded with the fluorescent probe calcein-AM and exposed to an osmotic challenge of different osmolytes during live imaging. Exposure of the tissue to aCSF containing an additional 100 mOsm led to abrupt shrinkage of the choroid plexus tissue when the osmotic challenge was generated by addition of mannitol (−0.9 ± 0.1% s^−1^, n = 5, P <0.001) or NaCl (−0.7 ± 0.0% s^−1^, n = 5, P < 0.001, Fig. 7a). In contrast, a 100 mOsm hyperosmotic challenge with KCl as the osmolyte induced instant swelling of the choroid plexus (0.4 ± 0.1% s^−1^, n = 5, P < 0.01), with no detectable delay (Fig. 7a). Application of the NKCC1 inhibitor bumetanide abolished the KCl-mediated volume increase of the choroid plexus, which then responded to the osmotic challenge with a shrinkage (−0.8 ± 0.1 % s^−1^, n = 5, P < 0.001) comparable to that obtained with the two other osmolytes (Fig. 7a). With the elevated [K^+^]_o_, the NKCC1 became inwardly directed and transported its solutes and water into the choroid plexus tissue, thereby accumulating intracellular water even when the tissue faced an osmotic gradient promoting outward water flux. NKCC1-mediated water transport can thus promote fluid movement in choroid plexus independently - and even against – a bulk osmotic gradient.

**Fig. 7.**
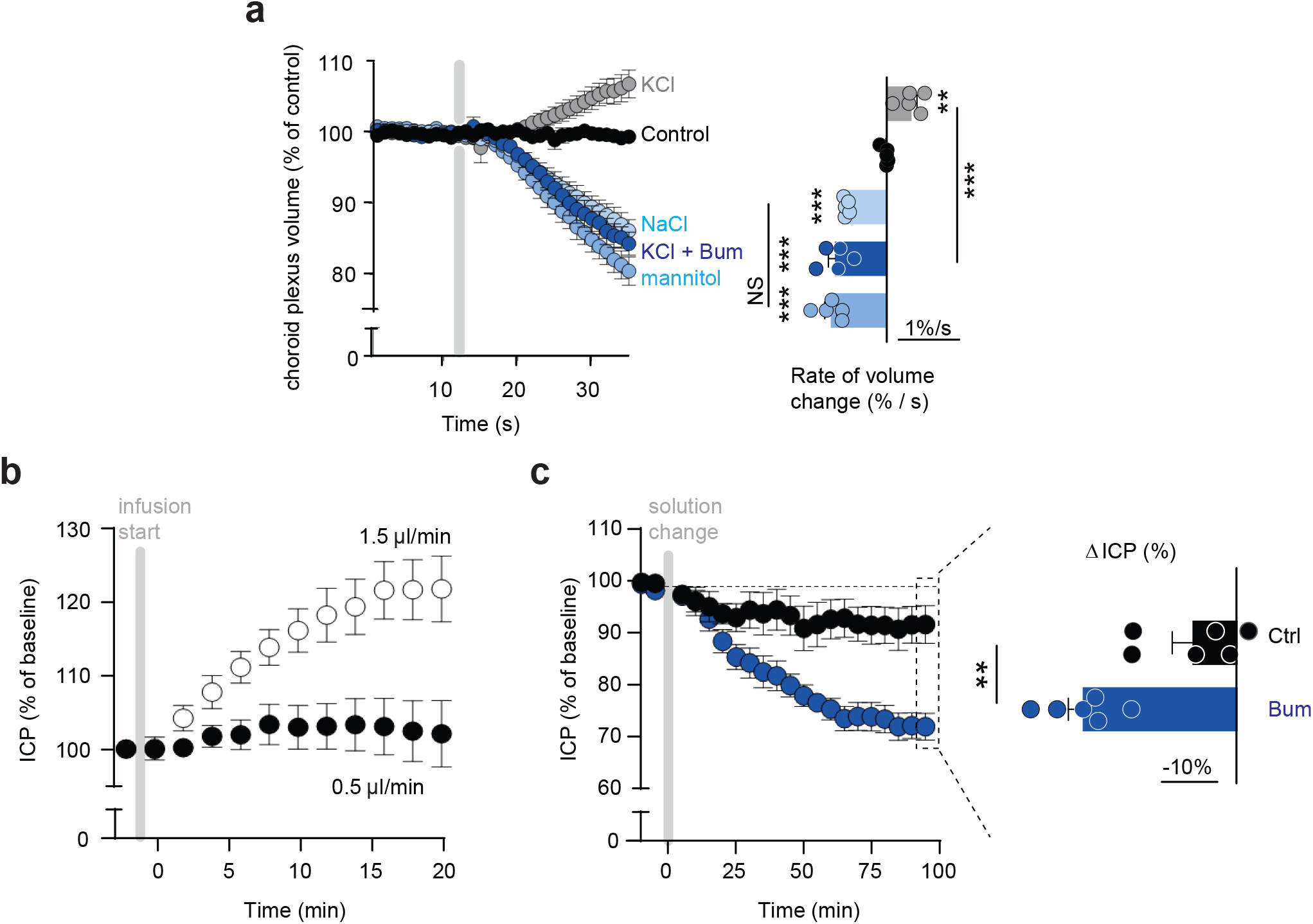
NKCC1-mediated fluid flow and contribution to ICP. **a,** Calcein fluorescence in rat choroid plexus as a function of time with exposure to control aCSF or +100 mOsm aCSF adjusted with mannitol, NaCl, or KCl in the absence or presence of bumetanide, when indicated by grey box, n = 5. Summarized data illustrated in right panel with statistical significance tested with one-way ANOVA followed by Tukey’s multiple comparisons, **P < 0.01, ***P < 0.001, NS = not significant. **b,** ICP recordings during intraventricular infusion of aCSF at rates of 0.5 (n = 6) or 1.5 µl min^−1^(n = 4) over 20 min. Unpaired two-tailed *t*-test was applied to evaluate statistically significant differences at the end of the procedure. **c,** The intracranial pressure of rats treated with slow intraventricular infusion (0.5 μl min^−1^) ± bumetanide for 100 min. Controls are from **b**, n = 6. Data are presented as % of control ICP as a function of time with each dot representing the mean value over a 5 min period. The endpoints are demonstrated in the right panel with unpaired two-tailed *t*-test employed for statistical analysis, **P < 0.01.

### NKCC1 inhibition lowers ICP

To test whether the CSF secretion can directly modulate the ICP, we measured the ICP epidurally in anesthetized and ventilated rats while mimicking a hypersecretion of CSF. The basal ICP was 4.3 ± 0.4 mmHg, n = 10, prior to infusion of aCSF and remained stable during the experimental window with a slow infusion of aCSF (0.5 μl min^−1^) into one lateral ventricle (102 ± 5% of ICP prior to initiation of infusion, n = 6, P = 0.74, Fig. 7b). Mimicking a 25 % increase in basal CSF secretion by an enhanced infusion rate of aCSF (1.5 μl/min), we observed an elevated ICP (122 ± 4 % of ICP prior to initiation of infusion, n = 4, P < 0.01, Fig. 7b). This demonstrates that mimicked CSF hypersecretion directly affected the ICP. To test if reduction of CSF secretion, likewise, could modulate the ICP, we infused bumetanide-containing aCSF into the lateral ventricle of anesthetized and ventilated rats while measuring the ICP. Infusion of vehicle-containing aCSF (0.5 µl min^−1^) did not significantly change the ICP over the 100 min duration of the experiment (11 ± 4% reduction, n = 6, P = 0.1), whereas infusion of bumetanide-containing aCSF led to a pronounced decrease in the ICP (of 29 ± 2%, n = 6, P < 0.01, Fig. 7c). These findings provide a proof-of-principle that pharmacological inhibition of CSF secretion can indeed promote a decline in the ICP.

## Discussion

Here, we present evidence suggesting that the CSF production is not driven primarily by conventional osmosis. We observed a low trans-epithelial water permeability of the rat choroid plexus *in vivo*, no detectable osmotic gradient across the tissue, minor contribution from local inter-microvillar standing gradients, and a unique capability to transport fluid against osmotic forces in the bulk solution. Instead, of conventional osmosis, we propose that CSF, at least in part, is formed by transporter-mediated water transport, which is geared to function in the absence of a trans-epithelial bulk osmotic gradient. The cotransporters NKCC1, NBCe2, and the Na^+^/K^+^-ATPase in the luminal membrane of choroid plexus appeared as key contributors to CSF secretion in rats. By abolishing the NKCC1-mediated fraction of the CSF secretion, we obtained experimental proof-of-concept that this molecular target could serve to control and reduce the ICP in rats; this may constitute a first step towards a needed efficient and targeted treatment option for patients with disturbed brain fluid dynamics and elevated ICP.

CSF secretion is generally assumed to be entirely osmotic: water is driven from the blood into the ventricles by a sufficiently large osmotic gradient across a sufficiently water-permeable epithelium. The presence of AQP1 in the luminal membrane has served as support for this model ^62,63^. Genetic deletion of AQP1, however, reduced the CSF production by only 20% in mice ^62^, part of which could originate from the 80% drop in the central venous pressure of these mice upon systemic AQP1 deletion ^62^. In alignment with the lack of substantial AQP expression in the basolateral membrane of choroid plexus, we obtained a low trans-epithelial osmotic water permeability of the rat choroid plexus (estimated L_p_ = 9 × 10^−5^ cm s^−1^ Osm^−1^ per apparent choroid plexus area, similar to that observed in rabbit choroid plexus ^64^). The choroidal trans-epithelial osmotic water permeability compares to other epithelial tissues with low water permeability (e.g. the dog urinary bladder with L_p_ = 2 × 10^−5^ cm s^−1^ Osm^−1 65^) and is ~50-fold lower than epithelial tissues with well-established high water permeability (e.g. the rat kidney proximal tubules with L_p_ = 4 ×10^−3^ cm s^−1^ Osm^−1 66^). Accordingly, we found that the secretory process of CSF formation was relatively insensitive to the osmolality of the ventricular fluid, with each extra mOsm leading to only a 0.4% increase in the CSF production. With this low capacity for passive water permeability in the choroid plexus, a favorable osmotic gradient of approximately 280 mOsm would be required to sustain the observed CSF production rate at □7 μl min, However, no such osmotic gradients were observed in our experimental rats and pigs, nor in humans (this study and ^67^). Other animal studies report a slightly increased ventricular hyperosmolality of ∼5 mOsm ^68–71^. This discrepancy may arise from experimental or technical differences such as a prolonged window between animal death and the sample extraction, pre-treatment of animals with saline before sampling ^69,70^, or artifacts from improper pre-analytical handling (e.g., freezing of CSF and/or ruptured blood cells in the centrifuged CSF).

Taken together, these findings indicate that conventional osmosis is a negligible driving force in the CSF secretion. More notably, robust CSF secretion occurred even when a bulk osmotic gradient favoring fluid flow *from* the ventricles *to* the blood stream was imposed across the choroid plexus. CSF secretion occurred against gradients as large as ∼ 10 times higher than the blood pressure. Such continued fluid secretion in the face of an oppositely-directed bulk osmotic gradient has previously been observed, but not explained, in different animal species, i.e. cats and goats, and with different osmolytes ^72–75^. This observation suggests that a mode of water transport different from conventional osmosis must be employed to sustain the fluid flux across the choroid plexus epithelium.

Various models of how trans-epithelial isotonic fluid transport may occur in the absence of measurable osmotic gradients have been proposed over the years, as comprehensively reviewed by Damkier and colleagues ^14^. Some of these models rely on water and ion permeability of the tight junctions supporting the isotonic fluid transport ^14^. With the tight junctions encompassing only a slight fraction of the epithelial membrane, these junctions must have excessively large unit water permeability to sustain the measured fluid flow ^76^. With such high unit water permeability, the anticipated size of these pores is difficult to reconcile with their impermeability to the electrolytes required to generate the osmotic gradient. The resulting low, or non-existing, reflection coefficient could represent a challenge for the model of tight junction-based osmotic water transport ^76^. Other models rely on local osmotic gradients arising in flow-protected compartments within the basolateral spaces between epithelial cells ^14^. Their existence was proposed ^28,77^ but remains to be experimentally demonstrated ^78,79^. CSF secretion across the choroid plexus takes place from the basolateral to the luminal side with the junctional complexes located at the luminal border and the luminal membrane covered with (1-3 μm) microvilli (this study and ^80,81^, some terminating in a bulbous structure (this study and ^82^). This reversed anatomical organization provides poor support for a local build-up of osmolytes (standing gradients and unstirred layers ^28,77^) in intercellular diffusion-restricted spaces, since these appear on the opposite side of the choroidal epithelium ^14,19,76^. To evaluate if local osmotic gradients could arise between the microvilli of the choroid plexus and drive fluid flow, we performed mathematical modelling of this process. With the employed parameters, the model assigned <0.1% of the CSF secretion to local osmotic forces between the choroidal microvilli. Such lack of osmotic particle build-up near the membrane surface is supported by ion concentration measurements in the vicinity of the choroidal microvilli brush border with ion-sensitive microelectrodes ^83^ and comparison of the osmolality of newly-formed CSF collected at the choroidal surface with that of bulk-collected CSF ^84^.

Cotransporter-mediated water transport may thus be the missing link supporting CSF secretion in the absence of a trans-epithelial bulk solution osmotic gradient. NKCC1 and KCC belong to the growing number of cotransporters with the ability to mediate cotransporter-mediated water transport ^18,85,86^, for review, see ^21,87^. Such transporter-mediated ability to move water across the choroid plexus luminal membrane, independently of the bulk osmotic gradient, has been detected in mouse ^15^, rat (this study), and salamander (Zeuthen 1994) choroid plexus, the latter of which was assigned to KCC activity, rather than the NKCC1 observed in the rodents. This mode of water transport is observed with transporter-mediated substrate translocation alongside concomitant movement of a number of water molecules; a number that is independent of the osmotic gradient in the surrounding bulk solutions ^88^. Cotransporter-mediated water transport thus appears to take place via a process within the protein itself ^19,20,88^, the precise mechanisms of which remains elusive. A recent cryo-EM study of NKCC1 revealed 86 water molecules residing in its translocation pathway at the given conformation depicted in the structure ^89^. This number of water molecules is incompatible with the earlier proposed *occlusion model*^19,87,89^, which stipulated occlusion of the transported water molecules along with the transported substrates. However, such prominent number of water molecules in the substrate translocation pathway provides support for the working model of cotransporter-mediated water transport that is based on a *hyper-osmolar cavity.* In this model, the substrate(s) is held in a thermodynamically free state during its exit from its binding sites within the protein, and thereby builds up hyperosmolarity inside the exit cavity of the protein and subsequent fluid flow through the cotransporter protein (described in ^19,87^, simulated in ^20^, and modelled in ^90^). We anticipate future structural revelations of distinct conformational states of these transport proteins and subsequent molecular dynamics simulations thereof to provide mechanistic insight into the molecular mechanisms underlying cotransporter-mediated water transport.

NKCC1 and KCC1, but neither of the KCC2-4 isoforms, were expressed at the mRNA and protein level in choroid plexus of rats and pigs, in accordance with other studies ^15,91–93^, but conflicting with studies detecting KCC3 or KCC4 in the choroid plexus ^60,61^. With no functional KCC expression in the luminal membrane of rat, pig, or mouse choroid plexus, NKCC1 alone acted on the CSF-facing side of choroid plexus with an outwardly-directed K^+^ translocation (this study and ^15,33^). This unique transport direction is assigned to the high choroidal ion concentrations of Na^+^ and Cl^− 15^, which determines the electrochemical gradient and thus the ionic transport direction for a given protein. However, the NKCC1 transport direction may be reversed under certain circumstances such as during early development or in dissociated choroid plexus epithelial cells ^81,94,95^.

Detection and quantification of choroidal electrolyte transport mechanisms demonstrated the Na^+^/K^+^- ATPase catalytic subunit α1 in association with β1/β3 (and near-absence of the neuronal (α3) and glial (α2) isoforms) in addition to NKCC1 and NBCe2 amongst the highest expressed luminal electrolyte transport proteins in the choroid plexus, with lesser abundance of NHE1. To reveal the relative contribution of each of these Na^+^-coupled transporters to the CSF secretion across the luminal membrane, these transcriptional top hits of highly expressed electrolyte transporters were examined by *in vivo* determination of CSF secretion rates in rats by direct pharmacological inhibition. Quantification of their contribution to CSF secretion demonstrated robust contributions from the Na^+^/K^+^-ATPase, NKCC1, NBCe2, and none from NHE1, with absence of indirect effects arising from inhibitor-mediated cardiovascular changes. These findings were obtained with two different CSF secretion assays and underscored by the impact of these transporters on the intracellular [Na^+^] homeostasis in the choroid plexus epithelium. Although NHE1 did not contribute to CSF secretion, its expression in the luminal membrane of choroid plexus ^53^ was confirmed and its functional role in CSF pH dynamics demonstrated. Our data confirm earlier demonstration of the involvement of these transport proteins in CSF secretion ^15,58, 96–106^. However, the present data set demonstrates, for the first time, the relative contributions of these transport proteins to CSF secretion. Inhibitors of the different transport proteins are envisaged to reach their target transport proteins in all CSF-facing structures, i.e., the ependymal cell lining or the leptomeninges insofar as these transport proteins might be expressed there. Accordingly, the CSF secretion assays are not exclusive to choroidal CSF secretion, although this site of origin appears to dominate ^3^. Pharmacological approaches, as employed here, may be hampered by specificity issues, but they represent a clear advantage in the acuteness of their inhibitory action. Prolonged inhibition (or genetic deletion) of any of the implicated transport mechanisms may affect intracellular ion concentrations and/or pH. These predicted changes may indirectly affect neighbor transport mechanisms, and thus potentially add confounding elements to the experimental approach.

The different, but complementary, technical approaches employed in this study (ventriculo-cisternal perfusion, LI-COR, [Na^+^]_i_ measurements) produce experimental read-outs at distinct time scales (minutes to hours), and are based on both *in vitro* and *in vivo* experimentation, yet still produced similar results. Such alignment may facilitate drawing conclusions regarding the direct implication of the tested transport proteins in these processes.

With the delineation of the luminal membrane transport mechanisms supporting CSF secretion, it now remains to categorize the molecular pathways for water entry from the basolateral side of the choroid plexus. At this location reside the bicarbonate transporting mechanisms AE2 and NCBE ^107^, which rank as number 6 and 10 highest expressed choroidal plasma membrane transporters (of 233 transcripts) in the RNAseq data set obtained in this study, in contrast to the bicarbonate transporter NBCn1 with much lower abundance in the choroid plexus tissue (6 TPM). The importance of choroidal HCO_3_ transporters is evident from the ability of the carbonic anhydrase inhibitor acetazolamide to reduce CSF secretion (^3,105^, but their individual contributions remain elusive. Successful genetic deletion of select choroidal HCO_3_ transporters promoted a redistribution of other membrane transport mechanisms, which prevented firm conclusions of the CSF secretion capacity of the genetically deleted transporters.

Elevation of ICP is generally anticipated to occur following blockage of CSF efflux routes. While such manner of ICP elevation certainly accounts for the majority of hydrocephalus cases, CSF hypersecretion, or mimicking thereof, may cause the damaging brain fluid accumulation observed in connection with a range of different pathologies (this study and ^106^). Irrespective of the origin of the fluid accumulation, reduction of CSF secretion by pharmacological measures could be a future therapeutic route to lowering elevated ICP. Accordingly, acute inhibition from the intraventricular side of the choroidal transport mechanism NKCC1 implicated in CSF secretion reduced the ICP of healthy rats by 30%. Such decrease in bumetanide-mediated ICP reduction aligns well with this NKCC1 inhibitor preventing intraventricular hemorrhage-induced ventriculomegaly ^16^. With the human choroidal expression of NKCC1 ^109^, this protein - or another of the choroidal transport proteins involved in CSF secretion -or their potential upstream regulatory signaling pathways could serve as a molecular target for therapeutic treatment of certain conditions with disturbed CSF dynamics.

In conclusion, we here propose that CSF secretion across the choroid plexus does not occur by conventional osmosis, whereby electrolytes are transported across the epithelium with osmotically obliged water entering via parallel routes. A trans-choroidal osmotic gradient is absent and the water permeability of the choroid plexus epithelium thus too low to support the well-established rate of CSF secretion. Local osmotic gradients arising in the inter-microvillar space do not appear to exist and/or suffice to support the CSF secretion. The ability of choroid plexus to continue CSF secretion even when faced with unfavorable osmotic gradients thus promotes cotransporters, and their ability to support water transport independently of the prevailing osmotic gradient in bulk solutions, as the molecular mechanism supporting this pivotal fluid flow. Accordingly, we have identified the cotransporter NKCC1, the Na^+^/K^+^-ATPase, and the HCO_3_ transporter NBCe2 as key contributors to choroidal CSF secretion. Future research will, hopefully, reveal the molecular mechanisms underlying the cotransporter-mediated water transport, determine if the Na^+^/K^+^-ATPase and NBCe2 belong to the transport proteins capable of supporting such water flux, and quantify the contribution from different basolateral transport proteins sustaining the fluid movement from the interstitial space and into the choroid plexus epithelium. Such identification could promote the highly desired pharmacological targets accessible from the vascular compartment and aimed at reducing CSF secretion in the pathologies comprising brain fluid accumulation and elevated ICP.

## Abbreviations

aCSF: artificial cerebrospinal fluid
AQP: aquaporin
CSF: cerebrospinal fluid
CP: choroid plexus
ICP: intracranial pressure
i.p.: intraperitoneal
KCC: K^+^/Cl^−^ cotransporter
NBCe2: Na^+^-HCO_3_^−^ transporter
NHE: Na^+^/H^+^ exchanger
NKA: Na^+^/K^+^ ATPase
NKCC1: Na^+^/K^+^/2Cl^−^ cotransporter
NPH: normal pressure hydrocephalus
RT: room temperature
TPM: transcripts per million

## Acknowledgement

We thank Trine Lind Devantier, Anca Stoica, Rikke Lundorf, Carina Lynnerup Pedersen, Micael Loenstrup, Thomas Hartig Braunstein, Martin Fredensborg Rath (Faculty of Health and Medical Sciences, University of Copenhagen, Denmark), and Nina Rostgaard (Department of Neurosurgery, Rigshospitalet) for technical input and/or assistance. We thank Professor Thomas Jentsch (University of Hamburg, Germany) and Professor Jinwei Zhang (University of Dundee, UK) for providing us with the KCC1 and KCC4 antibodies and the team around Nikolaj Lilleør (Department of Cardiothoracic Surgery, Rigshospitalet, Denmark) for access to blood/CSF from experimental pigs.

## Funding

This work was funded by the Novo Nordisk Foundation; Tandem program (NNF17OC0024718, to NM, the Hartmann Foundation (to NM), the Friis’ Foundation (to ABS), the DFG (FOR2795 - Ro2327/13-1, to CRR), the Absalon foundation (to SGH and AHS), Toyota-Fonden (to SGH and AHS), Fidelity Bermuda Foundation (to VK), and Swiss National Science Foundation (182683, to VK).

## Authors’ contribution

N.M., E.K.O., A.B.S, D.B., K.T., T.L.T.B., C.R.R. designed the research; E.K.O., A.B.S., D.B., N.J.G., K.T., T.L.T.B, S.D.L., S.N.A., T.Z., S.G.H., A.H.S., F.V. executed experiments / analyzed data. P.R.K. and V.K. established the mathematical model. N.M., E.K.O., A.B.S. drafted the manuscript. All authors participated in the finalization of the manuscript.

## Competing interest

The authors declare that they have no competing interests.

## Consent for publication

Not applicable.

## Ethics approval

Animal experiments were in compliance with the Directive 2010/63/EU on the Protection of Animals used for Scientific Purposes. Use of human samples was approved by the Ethical Committees of the Capital Region of Denmark, approval number H-18046630.

## Notes

### Competing Interest Statement

The authors have declared no competing interest.

