## Supplemental figures for "Cerebrospinal fluid formation is controlled by membrane transporters to modulate intracranial pressure"

Suppl. Figure 1

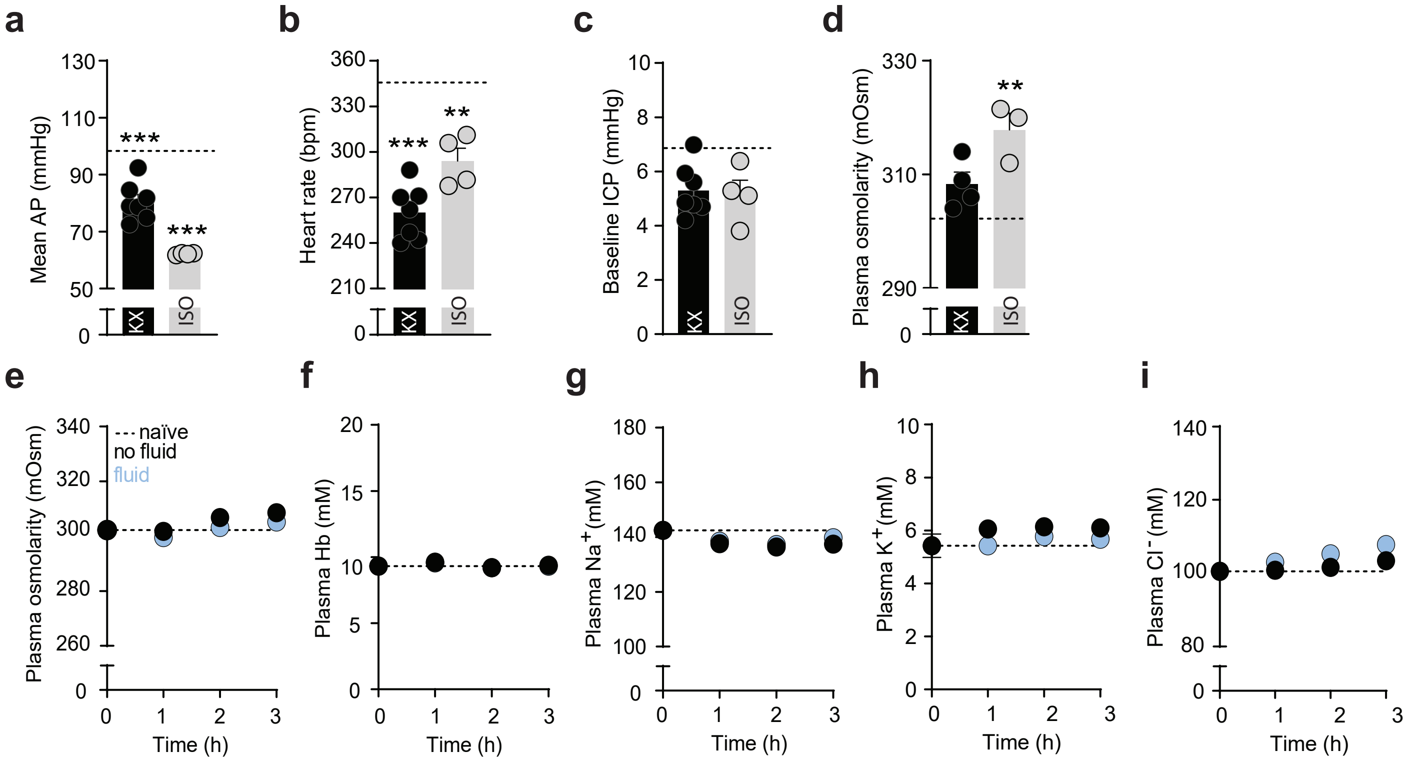


**Optimal physiological parameters under anesthesia.** To choose an anesthetic regiment optimal for retaining stable physiological parameters and thus correctly measured CSF secretion, rats were anesthetized with ketamine and xylazine (KX) or isoflurane (ISO) and ventilated over 3 h, while cardiovascular- and cranial measurements were recorded. Baseline values from non-anesthetized (NA) rats with telemetric pressure probes (ICP and MAP) or naïve rats, which were euthanized immediately after anesthesia induction, were included as dotted lines, n = 4. **a**, End mean arterial pressure (AP) measured from the femoral artery, KX: 82 ± 2 mmHg (n = 7), ISO: 62 ± 0 mmHg(n = 4), NA: 98 ± 2 mmHg (n = 4). **b**, End heart rate, KX: 255 ± 6 bpm (n = 7), ISO: 294 ± 8 bpm (n = 4), NA: 346 ± 8 bpm (n = 4). **c**, End intracranial pressure (ICP), KX: 5 ± 1 mmHg (n = 7), ISO: 5 ± 1 mmHg (n = 4), NA: 7 ± 3 mmHg (n = 4). **d**, End plasma osmolarity of KX-anesthesized rats (308 ± 2 mOsm) n = 4, ISO-anesthesized rats (318 ± 3 mOsm), n = 3, naïve rats (302 ± 2 mOsm) n = 4. Statistical significance from NA rats or from naïve rats was determined by one-way ANOVA with Dunnett's multiple comparisons *post-hoc* test. ***P < 0.001, **P < 0.01. We chose to anesthetize all ventilated rats with KX rather than ISO, since KX ensured stable plasma osmolarity and better mean arterial blood pressure compared with ISO. We tested whether hydration (5 ml subcutaneous 0.9 % isotonic saline, distributed 4 different places) was beneficial to cardiovascular parameters of rats during long anesthetic protocols using KX (n = 5). Control rats had no fluid treatment (n = 5). Baseline values were from naïve rats euthanized immediately after anesthesia induction (n = 4). **e**, Plasma osmolarity was stable during the anesthetic protocol (1 h vs. 3 h; 302 ± 2 vs. 309 ± 3 mOsm, P > 0.05, not different from naïve: 302 ± 2 mOsm, P > 0.05) and unaffected by fluid administration after ended experiment (no fluid vs. fluid; 309 ± 3 vs. 305 ± 3 mOsm, P > 0.05). **f**, Plasma Hb level was stable over 3 h (1 h vs. 3 h: 9.3 ± 0.4 vs. 9.4 ± 0.3 mM Hb, P > 0.05, which was not different from naïve: 9.3 ± 0.4 mM Hb, P > 0.05) and unaffected by fluid administration at termination (no fluid vs. fluid: 9.4 ± 0.3 vs 9.3 ± 0.3 mM Hb, P > 0.05). **g**, Plasma [$Na^+$] was stable over 3 h (1 h vs. 3 h: 143 ± 0 vs. 137 ± 2 mM $Na^+$, P > 0.05, which was not different from naïve: 143 ± 1 mM, P > 0.05) and unchanged by fluid treatment (no fluid vs. fluid: 137 ± 2 vs. 140 ± 0 mM $Na^+$, P > 0.05). **h**, Plasma [$K^+$] was stable over 3 h (1 h vs. 3 h: 5.4 ± 0.4 vs. 6.1 ± 0.2 mM $K^+$, P > 0.05, which was not different from naïve: 5.4 ± 0.4, P > 0.05) and unchanged by administration of fluid (no fluid vs. fluid: 6.1 ± 0.2 vs. 5.7 ± 0.1 mM $K^+$, P > 0.05). **i**, [$Cl^-$] concentration in plasma was stable over 3 h (1 h vs. 3 h: 101 ± 1 vs. 103 ± 1 mM $Cl^-$, P > 0.05, which was not significantly different from naïve: 101 ± 1 mM, P > 0.05) and unaffected by fluid administration (no fluid vs. fluid: 103 ± 1 vs. 108 ± 0 mM $Cl^-$, P > 0.05). Statistical significance was determined with two-way ANOVA with Tukey's multiple comparison *post-hoc* test. Because fluid administration was not beneficial to the tested physiological parameters, we evaluated that perioperative fluid administration was redundant during terminal experimental procedures.

Suppl. Figure 2

**a**

| Acc. number | Target | Sequence |
|---|---|---|
| NM_031798.2 | NKCC1 | *F:* GGC AAG ACT TCA ACT CAG CCA<br>*R:* TCA ACA AGG TCA AAC CTC CAT CA |
| NM_019229.2 | KCC1 | *F:* CGA GCG GAC ACT GAT GAT GG<br>*R:* GCG CCT GTC ACA GAC TCA TC C |
| NM_0013936675 | KCC2 | *F:* GAT GCA CGA GAG CGA CAT CT<br>*R:* CCG AGT GTT GGC TGG ATT CT |
| NM_001109630.1 | KCC3 | *F:* CAA GTC CCA CGA AGC AAA GC<br>*R:* AAA GGT CGG ATC ACG CAG AG |
| NM_001013144.2 | KCC4 | *F:* CCA GGA TGC TCA ACT CGT CC<br>*R:* CTG TTC AGC CCT TCG GTC AA |
| NM_012504.1 | ATP1a1 | *F:* GCT TTC CTG TCC TAC TGC CC<br>*R:* TTC CGC ACC TCG TCA TAC AC |
| NM_012505.2 | ATP1a2 | *F:* TCC CTT GGA GAC CCG CAA TA<br>*R:* CAT GGC TAT CGG CGT CTG TC |
| NM_012506.1 | ATP1a3 | *F:* TGG CCG AGA TTC CCT TCA AC<br>*R:* ATC GGT TGT CAT TGG GGT CC |
| NM_013113.2 | ATP1b1 | *F:* AAG AAG GAG TTT TTG GGC AGG A<br>*R:* CAC TTG GAT GGT CCC GAT GAA |
| NM_012507.3 | ATP1b2 | *F:* CAG GTG GTT GAG GAG TGG AAG<br>*R:* TCG ATC CTG GTA CTT GGG GG |
| NM_012913.1 | ATP1b3 | *F:* CAT CTA CAA CCC GAC CAG CG<br>*R:* TGT GAA GAG TGC AGC CAA GAA |
| NM_212512.2 | NBCe2 | *F:* ACC ACT GTC AAC GAC TCA GC<br>*R:* ATG AAG GAC ATG AGG GCC AAG |
| NM_012652.2 | NHE1 | *F:* AAC GGC TGC GGT CCT ATA AC<br>*R:* GGT TCA TAG GCC AGT GGG TC |
| NM_031144.3 | Actb | *F:* ACA ACC TTC TTG CAG CTC CTC<br>*R:* CTG ACC CAT ACC CAC CAT CAC |
| NM_213557.1 | Rps18 | *F:* CCT GCG AGT ACT CAA CAC CAA<br>*R:* TGG TGA GGT CAA TGT CTG CTT T |

**Primer sequences for qPCR.** Accession number, target protein name, and primer sequences for qPCR. F: Forward, R: Reverse.

Suppl. Figure 3

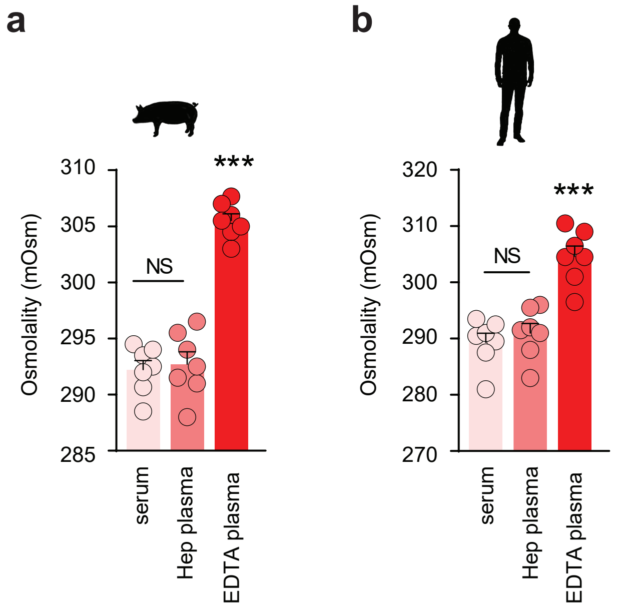


**Osmolarity in blood samples.** Osmolality of venous blood collected in heparin coated tubes (Hep plasma), EDTA coated tubes (EDTA plasma), or tubes without coating (plasma/serum) sampled from pigs (**a**) or healthy humans (**b**), n = 7 for both. EDTA coating increases the blood osmolality in pigs (plasma: 292 ± 1 mOsm, Hep: 293 ± 1 mOsm, EDTA 306 ± 1 mOsm) and humans (serum: 289 ± 2 mOsm, Hep: 291 ± 2 mOsm, EDTA: 305 ± 2 mOsm). One-way ANOVA with Tukey's multiple comparisons *post hoc* test, ***$P < 0.001$, NS = not significant. Blood osmolality was thus obtained from samples collected in heparin coated tubes for rats and pigs and tubes without coating (serum) in human.

Suppl. Figure 4

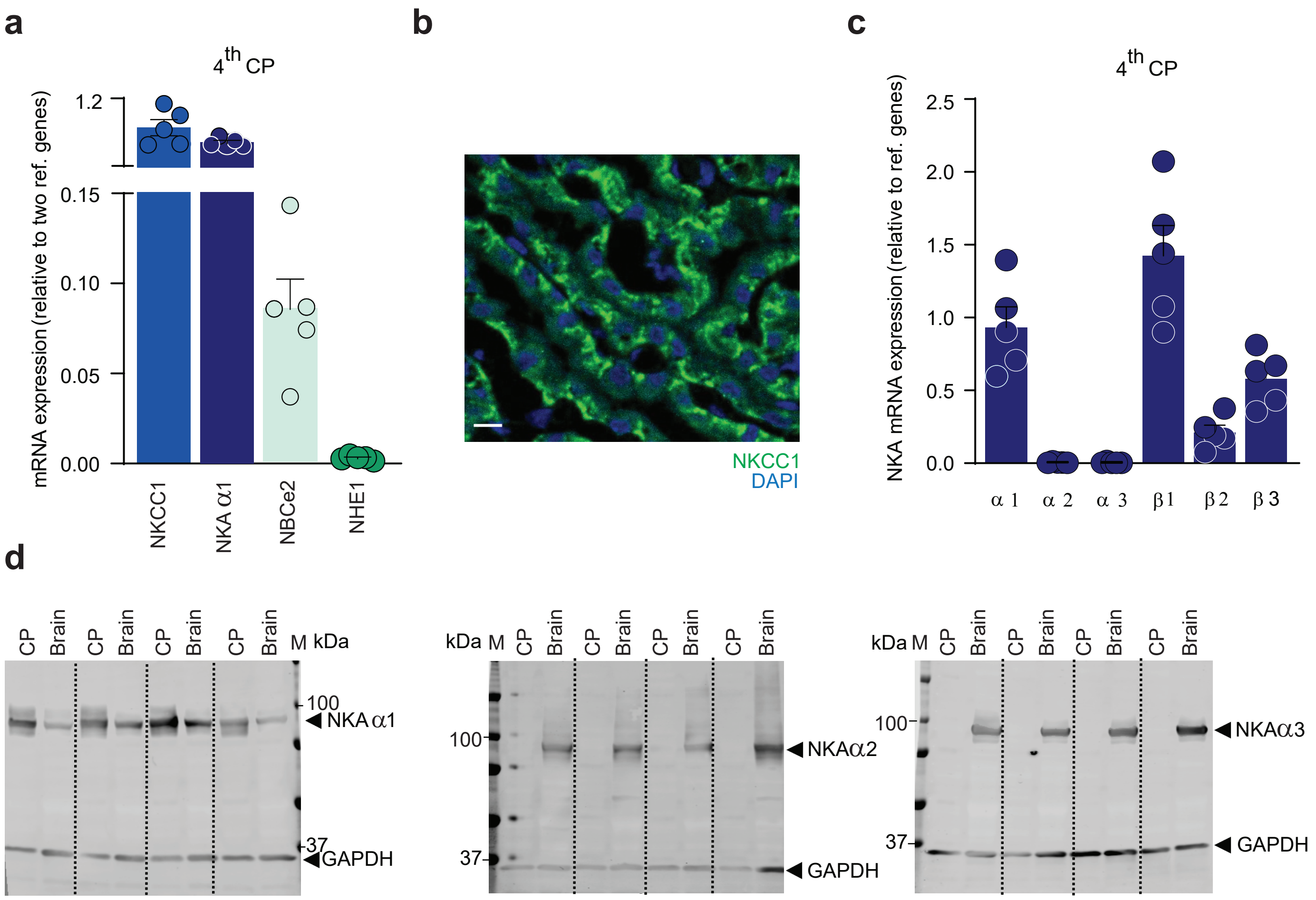


**Choroidal expression of transporters implicated in CSF secretion. a,** mRNA expression of NKCC1, $Na^+/K^+$-ATPase subunit α1, NBCe2, and NHE1 in choroid plexus from the 4$^{th}$ ventricle relative to two household genes determined from qPCR, n = 5. **b,** Representative immunostaining image of NKCC1 (green) in the CSF-facing membrane of the lateral choroid plexus from rat with DAPI-stained nuclei in blue. Scalebar 10 µm. **c,** mRNA expression levels in rat 4$^{th}$ choroid plexus of $Na^+/K^+$-ATPase subunit α1, α2, α3, $\beta$1, $\beta$2, and $\beta$3 relative to reference genes, n = 5. **d**, Original western blots of $Na^+/K^+$-ATPase subunit α1, α2, and α3 in lateral choroid plexus or whole brain tissue, n = 4. GAPDH was used as loading control.

Suppl. Figure 5

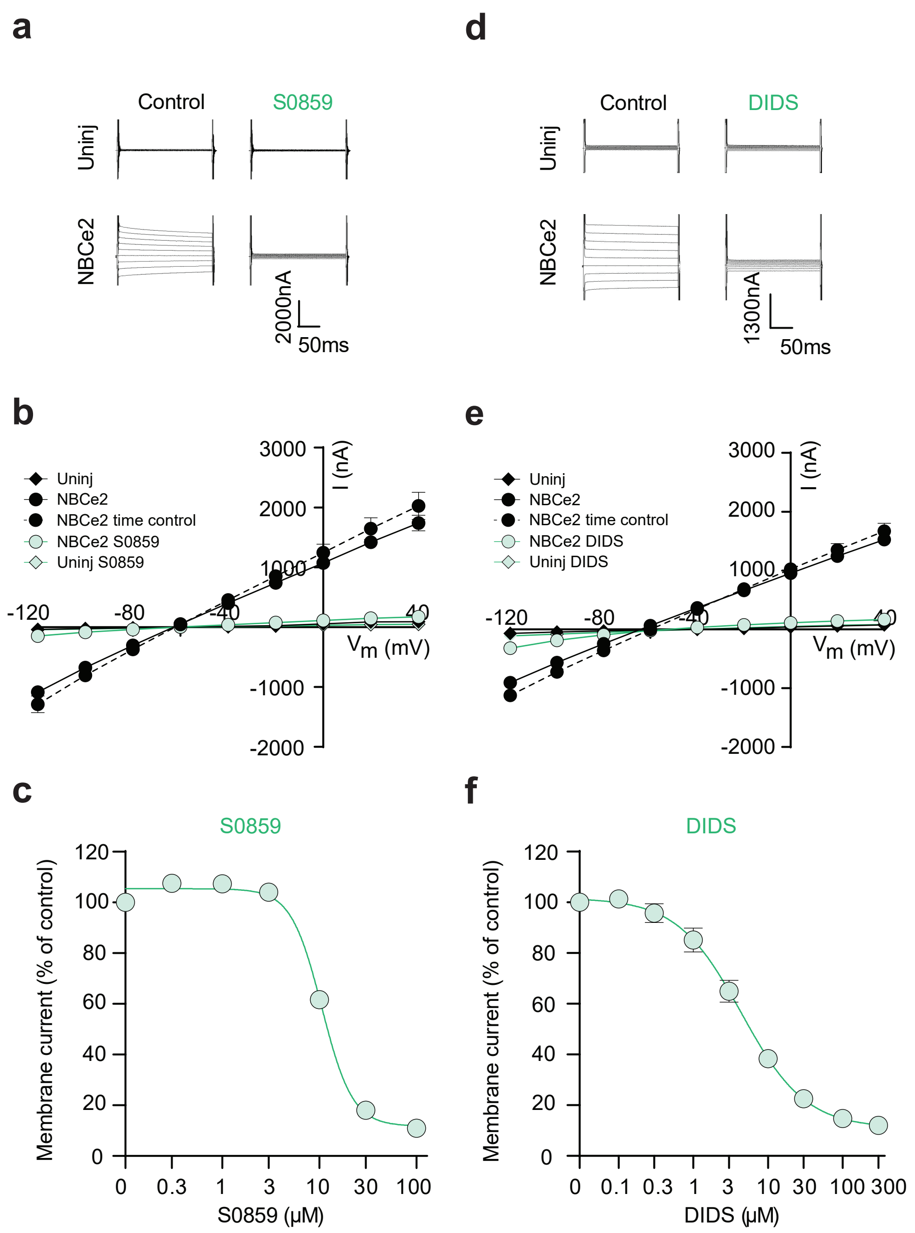


**Inhibitor potency of S0859 and DIDS on NCBe2. a,** Raw traces demonstrating membrane currents of uninjected *Xenopus leavis* oocytes (top panels) and NBCe2-expressing oocytes (lower panels) in control solution (left panels) or upon exposure to 100 µM S0859 (right panels). Membrane currents were recorded by application of 200 ms voltage steps from −120 mV to +40 mV in 20 mV increments from a holding potential of -60 mV. **b**, Summarized I/V relationship of uninjected and NBCe2-expressing oocytes (control and S0859) in addition to a time-control experiment to demonstrate stable NBCe2-mediated currents over time with no inhibitor included, n = 9 in each group. **c**, Dose-response curve of S0859 on the NCBe2-mediated membrane currents obtained in NBCe2-expressing oocytes presented as % of control current (obtained at $V_m$ = +40 mV) upon exposure to the different S0859 concentrations, n = 9. Error bars are within symbols. **d,** Raw traces demonstrating membrane currents of uninjected *Xenopus leavis* oocytes (top panels) and NBCe2-expressing oocytes (lower panels) in control solution (left panels) or upon exposure to 300 µM DIDS (right panels). Membrane currents were recorded by application of 200 ms voltage steps from −120 mV to +40 mV in 20 mV increments from a holding potential of -60 mV. **e**, Summarized I/V relationship of uninjected and NBCe2-expressing oocytes (control and DIDS) in addition to a time-control experiment to demonstrate stable NBCe2-mediated currents over time with no inhibitor included, n = 6 of each. **f**, Dose-response curve of S0859 on the NCBe2-mediated membrane currents obtained in NBCe2-expressing oocytes presented as % of control current (obtained at $V_m$ = +40 mV) upon exposure to the different DIDS concentrations, n = 6. Error bars are within symbols when not visible.

Suppl. Figure 6

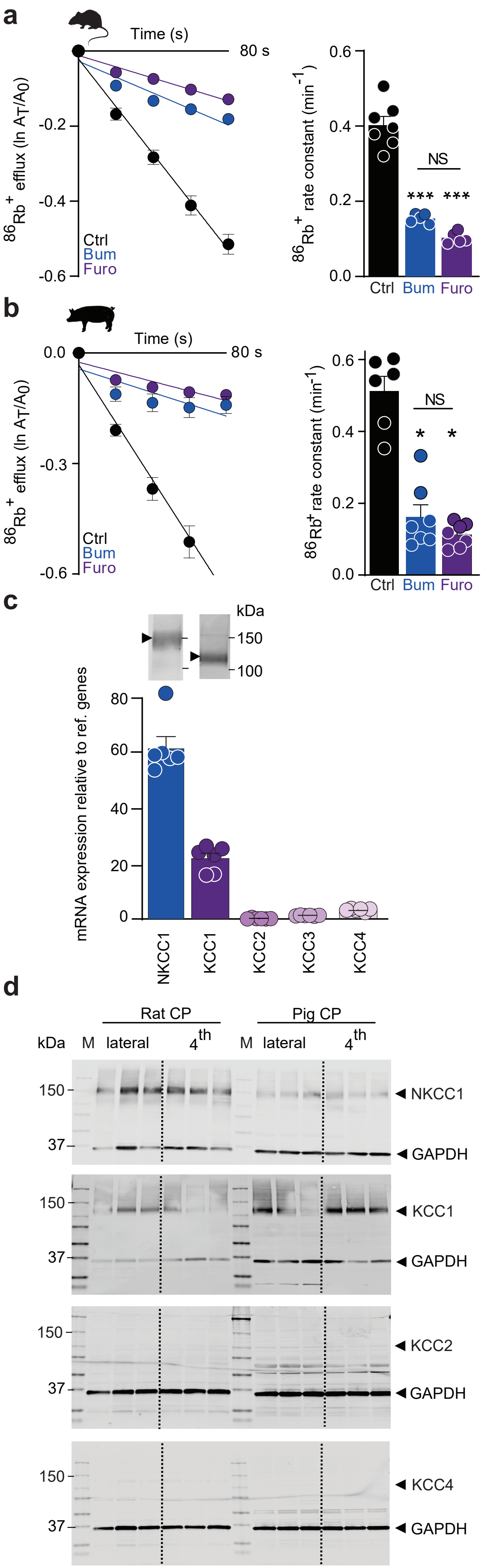


**Cation-chloride-coupled transporters in choroid plexus. a,** $^{86}Rb^{+}$ efflux from rat choroid plexus over time in control conditions (incl. vehicle, n = 7) and in the presence of bumetanide (20 µM, n = 5) or furosemide (1 mM, n = 5). Right panel; quantification of $^{86}Rb^{+}$ efflux rate constants (ctrl: 0.51 ± 0.04 $min^{-1}$, bum: 0.16 ± 0.03 $min^{-1}$, furo: 0.11 ± 0.01 $min^{-1}$) determined from linear regression analysis. Statistical significance determined with one-way ANOVA followed by Tukey's multiple comparisons *post-hoc* test. Asterisks above the bars indicate statistical difference from control conditions, ***; P <0.001, NS; not significant. . **b,** $^{86}Rb^{+}$ efflux from pig choroid plexus over time in control conditions (incl. vehicle, n = 6) and in the presence of bumetanide (n = 7) or furosemide (n = 7). Right panel; quantification of $^{86}Rb^{+}$ efflux rate constants (ctrl: 0.51 ± 0.04 $min^{-1}$, bum: 0.16 ± 0.03 $min^{-1}$, furo: 0.11 ± 0.01 $min^{-1}$) determined from linear regression analysis. Statistical significance determined with one-way ANOVA followed by Tukey's multiple comparisons *post-hoc* test. Asterisks above the bars indicate statistical difference from control conditions, *; P <0.05, NS; not significant. **c**, mRNA expression levels of NKCC1 and KCC1-4 isoforms in rat lateral choroid plexus by qPCR, relative to reference genes, see Methods, n = 5. Inset: Representative western blots of the expression of NKCC1 and KCC1 in rat choroid plexus. **d,** Representative western blots of NKCC1, KCC1, 2 and 4 in choroid plexus (lateral + 4$^{th}$) in rats and pigs, n = 3. The specificity and activity of all antibodies were previously verified (Steffensen et al, 2018). GAPDH was included as loading control.
