## supplementary data for "Cerebrospinal fluid formation is controlled by membrane transporters to modulate intracranial pressure"

Fluid velocity and solute concentration distribution along the longitudinal axis of a functional unit (FU) were determined through the set of coupled steady-state differential Equations S1 and S2 originally proposed by Diamond & Bossert [28] with the boundary conditions specified in Equations S3 to S5:

$$\frac{4\varphi(x)}{\rho \cdot d} + D\frac{d^2C}{dx^2} - C(x)\frac{dv}{dx} - v(x)\frac{dC}{dx} = 0, \tag{S1}$$

$$\frac{dv}{dx} = \frac{4L_\mathrm{p}}{d}[C(x) - C_0] \tag{S2}$$

$$C(l) = C_0 \tag{S3}$$

$$v(0) = 0 \tag{S4}$$

$$\left.\frac{dC}{dx}\right|_{x=0} = 0 \tag{S5}$$

Here, $\varphi$ is the local solute flux, $\rho$ is the cerebrospinal fluid (CSF) density, $d$ is the FU diameter, and $x$ is the location along the longitudinal axis of the FU, starting from 0 at the surface of the luminal cell membrane and ending at $l$, the length of a microvillus. $D$ is the diffusion coefficient, $C$ is the solute concentration, $v$ is the fluid velocity, $L_\mathrm{p}$ is the permeability of the luminal epithelial membrane, and $C_0$ is the bulk solute concentration in the ventricular CSF. Solute flux through the base of the FU is accounted for by a correspondingly increased flux from the side of the FU within the first $d/(2 \cdot l)$ of the functional unit length.

The diameter, $d$, of a functional unit (FU) is obtained by calculating the equivalent hydraulic diameter of the volume spanned by four neighboring microvilli, namely

$$d = \frac{4A_\mathrm{CS}}{P_\mathrm{CS}} = \frac{p^2 - \pi r^2}{p + \left(\frac{\pi}{2} - 2\right)r} \tag{S6}$$

Here, $A_{CS}$ and $P_{CS}$ represent the FU cross-sectional area and perimeter, respectively. The pitch, $p$, is the separation distance of adjacent microvilli, and $r$ is the radius of a microvillus. The pitch and number of functional units, $N$, are determined as

$$p = \sqrt{\frac{1}{n}} \tag{S7}$$

$$N = n \cdot A_{\mathrm{app}} \tag{S8}$$

where $n$ is the number of microvilli per unit area, $A_{\mathrm{app}}$ is the apparent luminal surface membrane area, i.e., without accounting for the surface extension by microvilli. The folding factor, $FF$, is defined as the ratio of the actual to the apparent luminal membrane surface area with

$$FF = A_{\mathrm{act}}/A_{\mathrm{meas}} \tag{S9}$$

$$A_{\mathrm{act}} = N(p^2 + 2\pi r l) \tag{S10}$$

The rate of solute removal from the ventricular space by bulk flow of CSF, Φ, is calculated as

$$\Phi = V_p \cdot \rho \cdot C_0 \tag{S11}$$

where $V_p$ is the measured CSF secretion rate, $\rho$ is CSF density, and $C_0$ is the bulk solute concentration. This solute removal rate is assumed to be equal to the solute transfer rate into all FUs. The solute flux, φ, through the sides and bottoms of all FUs is then

$$\varphi = \frac{\Phi}{N\left(\frac{\pi d^2}{4} + \pi d l\right)} \tag{S12}$$

The CSF production rate predicted by the model, $Q$, is obtained as

$$Q = N \cdot q \tag{S13}$$

$$q = \frac{\pi d^2}{4} v(l) \tag{S14}$$

where $q$ is the calculated rate of water release from one FU and $v(l)$ is the flow velocity at the end of the FU as provided by Equations S1 to S5. Parameter values used in the model are provided in Table S1.

**Table S1-** Parameter values employed in the osmotic water transfer model

| Model parameter (unit) | Symbol | Value | | Source |
|---|---|---|---|---|
| | | Derived | Measured | |
| Measured CSF secretion rate (µl/min) | $V_p$ | | 6.8 | This study |
| Microvillus length (µm) | $l$ | | 1.71 | This study |
| Microvillus radius (µm) | $r$ | | 0.059 | This study |
| Number of microvilli per unit area (1/µm$^2$) | $n$ | | 18 | This study |
| Choroid plexus apparent area (cm$^2$) | $A_{app}$ | | 4.6 | This study |
| Diffusion coefficient (cm$^2$/s) | $D$ | | $1.5 \cdot 10^{-5}$ | Ref. 110 |
| Luminal membrane permeability for $A_{act}$ (cm·s$^{-1}$·Osm$^{-1}$) | $L_p$ | | $1.4 \cdot 10^{-5}$ | This study |
| Bulk solute concentration in CSF (Osm) | $C_0$ | | 0.307 | This study |
| Density of CSF (g/ml) | $\rho$ | | 1.00 | Ref. 49 |
| Diameter of functional unit, FU (µm) | $d$ | 0.212 | | Eq. S6 |
| Microvilli separation distance, pitch (µm) | $p$ | 0.236 | | Eq. S7 |
| Number of functional units (-) | $N$ | $8.28 \cdot 10^{9}$ | | Eq. S8 |
| Folding factor (-) | $FF$ | 12.37 | | Eq. S9 |
| Choroid plexus actual area (cm$^2$) | $A_{act}$ | 56.91 | | Eq. S10 |
| Solute transfer rate (mmol/s) | $\Phi$ | $3.48 \cdot 10^{-5}$ | | Eq. S11 |
| Solute flux (mmol·s$^{-1}$·cm$^{-2}$) | $\varphi$ | $3.57 \cdot 10^{-7}$ | | Eq. S12 |
